# Mutational Analysis of an Antimalarial Drug Target *Pf*ATP4

**DOI:** 10.1101/2024.02.21.581445

**Authors:** Swaksha Rachuri, Binod Nepal, Anurag Shukla, Aarti Ramanathan, Joanne M. Morrisey, Thomas Daly, Michael W. Mather, Lawrence W. Bergman, Sandhya Kortagere, Akhil B. Vaidya

## Abstract

Among new antimalarials discovered over the past decade are multiple chemical scaffolds that target *Plasmodium falciparum* P-type ATPase (*Pf*ATP4). This essential protein is a Na^+^ pump responsible for the maintenance of Na^+^ homeostasis. *Pf*ATP4 belongs to the type 2D subfamily of P-type ATPases, for which no structures have been determined. To gain better insight into the structure/function relationship of this validated drug target, we generated a homology model of *Pf*ATP4 based on SERCA, a P2A-type ATPase, and refined the model using molecular dynamics in its explicit membrane environment. This model predicted several residues in *Pf*ATP4 critical for its function, as well as those that impart resistance to various *Pf*ATP4 inhibitors. To validate our model, we developed a genetic system involving merodiploid states of *Pf*ATP4 in which the endogenous gene was conditionally expressed, and the second allele was mutated to assess its effect on the parasite. Our model predicted residues involved in Na^+^ coordination as well as the phosphorylation cycle of *Pf*ATP4. Phenotypic characterization of these mutants involved assessment of parasite growth, localization of mutated *Pf*ATP4, response to treatment with known *Pf*ATP4 inhibitors, and evaluation of the downstream consequences of Na^+^ influx. Our results were consistent with modeled predictions of the essentiality of the critical residues. Additionally, our approach confirmed the phenotypic consequences of resistance-associated mutations as well as a potential structural basis for the fitness cost associated with some mutations. Taken together, our approach provides a means to explore the structure/function relationship of essential genes in haploid organisms.

**Significance Statement:** *Plasmodium falciparum* ATP4 (*Pf*ATP4) is a Na^+^ efflux pump and represents an important target for antimalarial drugs with nanomolar potency. However, the structure of *Pf*ATP4 is unknown, prompting the development of new methodologies to investigate the structure/function relationship. Here, we introduce a dynamic homology modeling approach to (a) identify key residues essential for *Pf*ATP4 function and (b) provide a structural basis to understand resistance-associated mutations. To validate these predictions, we developed a genetic system to manipulate the PfATP4 gene to assess the phenotypic consequences of such changes. Our results support the utility of combining homology modeling and genetics to gain functional insights into an antimalarial drug target.

## Introduction

As the continued threat of resistance to mainstay antimalarials grows in endemic regions, significant efforts have been undertaken to discover new antimalarial drugs. Among the many antimalarials discovered over the past decades are multiple chemical scaffolds targeting *Pf*ATP4, a *Plasmodium falciparum* Na^+^/H^+^ pump (1–6). Two of these compounds have progressed to Phase IIb clinical trials, showing potent activity with extremely rapid clearance of circulating *P. falciparum* and *P. vivax* parasites (7, 8). Inhibition of *Pf*ATP4 results in an influx of Na^+^ in the parasite cytoplasm followed by dramatic consequences, including swelling of the parasite (9), changes to the lipid composition of the parasite plasma membrane (PPM) (10), induction of premature schizogony (10), and eryptosis (3), all of which contribute to parasite death.

*Pf*ATP4 belongs to a superfamily of P-type ATPases that use ATP hydrolysis to transport ions or metabolites across membranes (11, 12). These pumps are divided into five classes, P1 to P5, and are further divided into subclasses (A to D) based on their function and membrane localization (12–14). *Pf*ATP4 belongs to the P2-type ATPases, of which the sarco/endoplasmic reticulum Ca^2+^ ATPase (SERCA) and Na^+^/K^+^ ATPase have been extensively investigated (14). *Pf*ATP4 is further classified into the P2D subfamily, which contains pumps in fungi, lower plants, and some unicellular eukaryotes (15–17). Pumps in this family transport monovalent cations, such as Na^+^, and are called the ENA (***e****xitus **na**trus*) ATPases (15–17). Phylogenetic analysis of P2D-type ATPases revealed distinct clades within this subclass, one comprising of pumps from fungi, kinetoplastids, bryophytes, and Entamoeba, and the other consisting of apicomplexan, chromerid, stramenophile, chlorophyte, and dinoflagellate pumps **(Fig. 1a)**.

**Figure 1:**
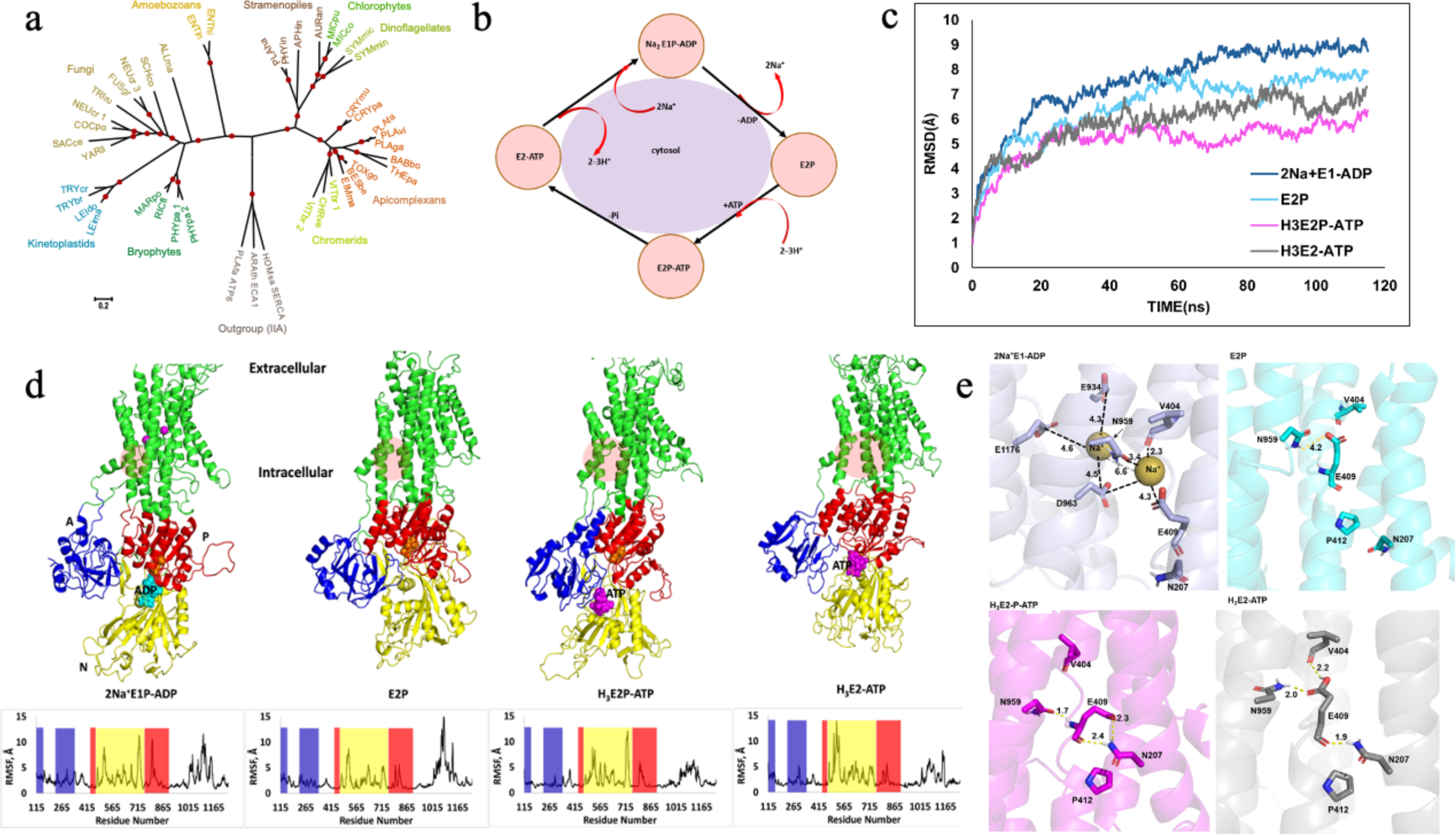
*Pf*ATP4 models using SERCA as a template. **(a)** Phylogenetic analysis of P2D-type ATPases, depicted as a radial tree diagram with P2A-type ATPases as outgroups; Red dots indicate >90% bootstrap support. A rectangular phylogram version of the tree is shown in **Fig. S1**. **(b)** Catalytic and intermediate states of PfATP4 shows the possibility of 4 conformational states namely 2Na^+^E1P-ADP, E2P, H_3_E2P-ATP and H_3_E2-ATP **(c)** trajectory traces from MD simulations of the 4 states shows all states are energetically stable from 25-115ns **(d)** three-dimensional molecular models of the 4 conformational states are shown as ribbons and colored according to their domain architecture (TM helices – green, A-domain – blue, N-domain – yellow and P-domain – red). The predicted PfATP4 inhibitor pocket is depicted as a pink circle in the TM domain. RMSF maps for each conformational state are shown below the structures, and A, N, and P domains are marked for clarity **(e)** Hydrogen bonding pattern at the proximity of the Na+ coordination site in each conformational state is shown with H-bonds marked as dotted yellow lines and labeled with distance, participating residues are shown as licorice sticks and colored atom type (carbon: grey/cyan/magenta, nitrogen: blue, oxygen: red) and labeled with residue number.

At present, no structure of a P2D-type ATPase has been determined. Since *Pf*ATP4 is a validated and attractive antimalarial drug target, it is crucial to understand the structure/function relationship of this protein. So far, attempts to express functional *Pf*ATP4 in heterologous systems have been unsuccessful, preventing such investigations. To circumvent this limitation, we developed a high-confidence homology model of *Pf*ATP4 undergoing its catalytic cycle in its explicit membrane environment. Using this model, we identified several critical residues likely involved in *Pf*ATP4 function. We performed mutational analysis of these residues using a genetic system that permits assessment of their consequences on parasite biology. In addition to identifying residues involved in *Pf*ATP4 function, our model was also used to assess the structural consequences of mutations conferring resistance to different *Pf*ATP4 inhibitors. In particular, we identified a possible structural basis for the fitness defect observed with the development of certain mutations. Mutations resulting in fitness defects can reduce the spread of resistance; therefore, it is crucial to study their structural consequences.

## Results

### Homology Modeling using SERCA Intermediates2

P-type ATPases cycle between four states, accompanied by massive structural rearrangements of the cytosolic and transmembrane (TM) domains **(Fig. 1b)** (13, 18–20). The structure of each state has been solved for SERCA, a P2A-type ATPase. Since no structures are available for *Pf*ATP4 or other P2D-type ATPases, we created a homology model and refined the structures for the four conformational states of *Pf*ATP4 in the Na^+^ transport cycle using the SERCA protein as a template. Analysis of the trajectories from molecular dynamics (MD) simulations showed that all systems are well equilibrated after 25 ns of simulation and have stable trajectories for the next 100 ns **(Fig. 1c)**. A comparison of these trajectories for the different conformational states reveals that the E2-ATP states form the most stable conformations. Mapping the root mean square fluctuations (RMSF) of the residues from different domains during the simulations showed that TM3, TM10, and the N-domain had major conformational changes in all the states while the E1 state additionally showed fluctuations in the P-domain **(Fig. 1d)**. In the modeled structures, E409, E934, D963, and E1176 formed the coordination sites for binding Na^+^; these residues align with E309, E771, D800 and E908 of SERCA that form the coordination sites for binding Ca^2+^ (21). Previous studies on the SERCA protein indicate that the E2 to E1 transition is triggered by the deprotonation of E309 (22) and a ∼110° rotation of the A-domain (23), leading to the large movement in TM1 and TM2. We analyzed the intramolecular interactions, especially H-bond interactions, in the vicinity of E409 and the ion binding sites in the four models **(Fig. 1e)**. In the 2Na^+^E1-ADP state, the two Na^+^ ions were stably coordinated at the ion binding sites throughout the simulation. The simulations also revealed that in addition to E409, E934, D963, and E1176 residues, the backbone carbonyl of V404 and the side chain of N959 also helped coordinate the Na^+^ ions in the 2Na^+^E1-ADP state. These two residues are also conserved in SERCA and participate in Ca^2+^ binding in the E1P state (24). The removal of the two Na^+^ ions led to the E2P state in which E409, E934, and E1176 are still deprotonated. The deprotonated E409 did not engage in an H-bond interaction with any residues (**Fig. 1e)**. In the H_3_E2P-ATP state, E409 is protonated, and it engaged in an H-bond interaction with N207 while the amine backbone of E409 engaged in an H-bond interaction with the N957 amine side chain. The dephosphorylated H_3_E2-ATP state had a significantly different H-bond interaction network than the other states. The protonated E409 engaged in three H-bond interactions: the side chain carbonyls H-bond with the backbone carbonyl of V404 and the N957 side chain, whereas the backbone carbonyl made an H-bond with the N207 side chain **(Fig. 1e)**. These findings indicate that the protonated E409 is crucial for stabilizing the H_3_E2-ATP state. An E309N single mutation in SERCA protein destabilized the E2 state and transitioned to an E1-like conformation (22). The loss of H-bonding between V203 and E309 was found to be a trigger for the E2 to E1 transition in SERCA (22). Besides the above intramolecular interactions, the study suggested that the A and N domains are more compact in the E2 state than in the E1 state (25). We measured the center of mass distances between the A and N domains in our *Pf*ATP4 models, which are presented in **Table S2**. The E2P state had the shortest distance (∼27Å), while the other 3 states had a ∼50Å distance separating the A and N domains. The displacement of ∼25Å from the E1 to E2 transition is similar to the SERCA protein E1 to E2 transition (26). The size of the binding cavity in each structure was calculated using CASTp (27) (Ver 3.0). The solvent-accessible volume of the binding cavity was the highest in H_3_E2-ATP (614Å^3^), followed by H_3_E2P-ATP (328Å^3^) and 2Na^+^E1P-ADP (292 Å^3^). The size of the binding cavity was the least in the E2P state (180 Å^3^).

AlphaFold (28) is an artificial intelligence program that can predict proteins’ three-dimensional structures from a given sequence using deep-learning neural network models. The equilibrated models of *Pf*ATP4 described above were compared with the AlphaFold-predicted structure, and the results are shown in **Table S2**. Superpositioning of the AlphaFold-generated *Pf*ATP4 model on the modeled 2Na^+^E1P-ADP, E2P, H_3_E2P-ATP, and H_3_E2-ATP structures resulted in root-mean-squared distance (RMSD) values of 6.24, 7.36, 7.31, and 6.96Å respectively for the complete structure and 3.29, 5.14, 3.72, and 3.09 Å, respectively, for the TM regions. These results suggest that the AlphaFold structure is not representative of any of the 4 conformational states of *Pf*ATP4.

### A genetic system to investigate *Pf*ATP4

We developed a genetic system to validate our homology model’s predictions and the importance of specific residues in *Pf*ATP4. First, we generated transgenic parasites expressing endogenous *Pf*ATP4 tagged with a c-Myc epitope at the C terminal end under the control of the TetR-DOZi aptamer system (29) using single crossover recombination in the NF54*attB* parasite line (called NF54*attB Pf*ATP4:3xMyc; **Fig. 2a**). In these parasites, expression of *Pf*ATP4 is maintained in the presence of anhydrotetracycline (aTc) (29). After successful integration, the knockdown of *Pf*ATP4 by the removal of aTc was confirmed using both anti-c-Myc and anti-*Pf*ATP4 antibodies **(Fig. 2b)**. As observed in a previous study (29), *Pf*ATP4 knockdown caused parasite demise within two cycles **(Fig. 2c)**. Western blot analysis of membrane preparations from these parasites revealed the presence of *Pf*ATP4 as a 140 kDa protein in SDS-polyacrylamide gel electrophoresis **(Fig. 2d)**.

**Figure 2:**
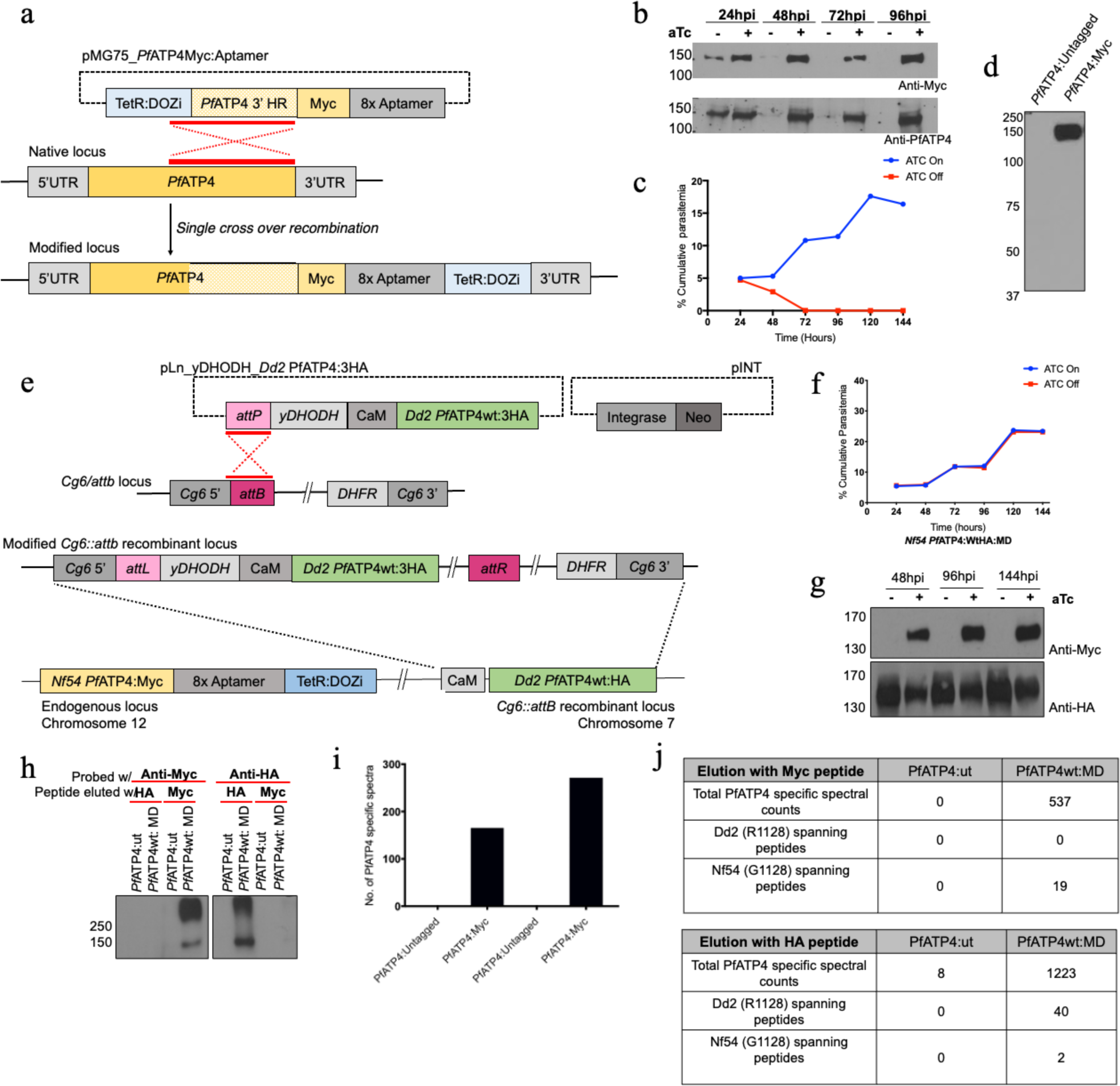
Genetic System to Investigate P*f*ATP4. **(a)** Schematic representation of the single crossover recombination strategy that was used to generate NF54*attB Pf*ATP4:3xMyc parasite line in which *Pf*ATP4 expression is regulated by the TetR-DOZi aptamer system at the endogenous locus. **(b)** Western blots of NF54*attB Pf*ATP4:3xMyc parasite lysate probed with anti-c-Myc and anti-PfATP4 antibody showing successful knockdown of endogenous PfATP4 upon removal of aTc within 48hpi. **(c)** Growth curves of NF54*attB Pf*ATP4:3xMyc parasites in the presence and absence of 250 nM aTc. Cultures were split 1:2 at 48 and 96 hours post invasion (hpi), and percent cumulative parasitemia was calculated by multiplying parasitemia and the split factor over time, representative of 3 biological replicates**. (d)** Western blots after SDS-PAGE (Panel 1) of membrane preparations from untagged and NF54*attB Pf*ATP4:3xMyc parasites probed with anti-c-Myc depicting a 140kd band corresponding to PfATP4. **(e)** Schematic representation of the merodiploid (MD) parasite line expressing aTc-regulated NF54attB PfATP4:3xMyc at the endogenous locus and Dd2 *Pf*ATP4:3xHA at the cg6∷attB locus expressed under the control of the calmodulin promoter. **(f)** Growth curves as estimated by cumulative parasitemia of wildtype and mutant merodiploid lines with and without aTc. Representative growth curves (of 3 biological replicates) of merodiploid *Pf*ATP4WT parasites. **(g)** SDS-PAGE followed by western blotting of merodiploid NF54*attB Pf*ATP4:3xMyc *Pf*ATP4(WT):3xHA parasite lysate maintained with and without aTc over 48, 96, and 144 hpi probed with anti-c-Myc and anti-HA to detect the expression of *Pf*ATP4 from the endogenous (top) and ectopic loci (bottom) indicating that the ectopic expression of *Pf*ATP4 remains stable upon knockdown of the endogenous PfATP4. **(h)** Western blots showing endogenous and ectopic *Pf*ATP4 expression from the merodiploid NF54*attB Pf*ATP4:3xMyc *Pf*ATP4(WT):3xHA parasites with anti c-myc and anti-HA showing co-dominant expression. **(i)** Proteomic analysis showing counts of peptide spectra originating from *Pf*ATP4 in two biological replicates of immunopulldowns from untagged and c-Myc-tagged parasites. Peptides common to both samples were considered as non-specific contaminants during pulldown and were eliminated during analysis. Except for *Pf*ATP4, no other proteins were represented uniquely in c-Myc-tagged pulldowns. **(j)** Proteomic analysis after immunopulldown (IP) of untagged (*Pf*ATP4:mut) and merodiploid (*Pf*ATP4:wtMD) parasites showing the presence of only cognate peptide spectra spanning the polymorphic region.

The NF54*attB Pf*ATP4:3xMyc parasite line was used to generate a merodiploid parasite line (NF54*attB Pf*ATP4:3xMyc/*Pf*ATP4_(mut)_:3xHA) in which a copy of constitutively expressed *Pf*ATP4 (originating from the Dd2 strain and tagged with a 3xHA epitope) was integrated at the cg6∷*attB* ectopic locus **(Fig. 2e)**. Conditional knockdown of endogenous *Pf*ATP4 expression by withdrawing aTc, was complemented by the presence of an ectopically expressed wild type *Pf*ATP4 **(Fig. 2f)**. Western blot analysis using anti-Myc, and anti-HA antibodies confirmed knockdown of endogenous *Pf*ATP4 and constitutive expression of ectopic *Pf*ATP4, respectively **(Fig. 2g)**. Immunopulldown experiments with either anti-c-Myc or anti-HA antibodies showed that the endogenous and ectopically expressed *Pf*ATP4 did not associate with each other **(Fig. 2h)**. This was further supported by the fact that a single polymorphism, G1128 in NF54 (the endogenous allele) and R1128 in Dd2 (the ectopic allele), distinguished the individual alleles. Proteomic analysis of eluates by LC-MS/MS revealed the presence of G1128 spanning peptides only in c-Myc eluted samples and R1128 predominantly in HA eluted samples **(Fig.2i-j)**. Together, these data validate our merodiploid system wherein the function of each allele of *Pf*ATP4 can be independently assessed.

### Assessing Likely Na^+^ Coordination Residues in *Pf*ATP4

As described above, our homology model identified residues E409, E934, D963, and E1176 as residues coordinating Na^+^ ions **(Fig. 3a)**. To validate these predictions, we conducted mutational analyses by creating multiple merodiploid lines with mutations in ectopic *Pf*ATP4 using the NF54*attB Pf*ATP4:3xMyc parasites **(Fig. 3b)**. Using these merodiploid lines, we assessed potential consequences to protein function and localization as well as to overall parasite growth. Of the four predicted Na^+^ coordination residues, we mutated E409 and E1176 to isoleucine residues to remove the charge and polarity while maintaining the relative size. The merodiploid parasite lines (*Pf*ATP4_E409I_, *Pf*ATP4_E1176I_) were grown in the presence and absence of aTc to assess whether the mutated allele could complement parasite growth. In the presence of aTc, the *Pf*ATP4 alleles from both loci were co-dominantly expressed **(Fig. S5b-c)**. Withdrawal of aTc from the medium resulted in cessation of parasite growth compared to complementation by *Pf*ATP4_WT_ **(Fig. 3-c-e)**.

**Figure 3:**
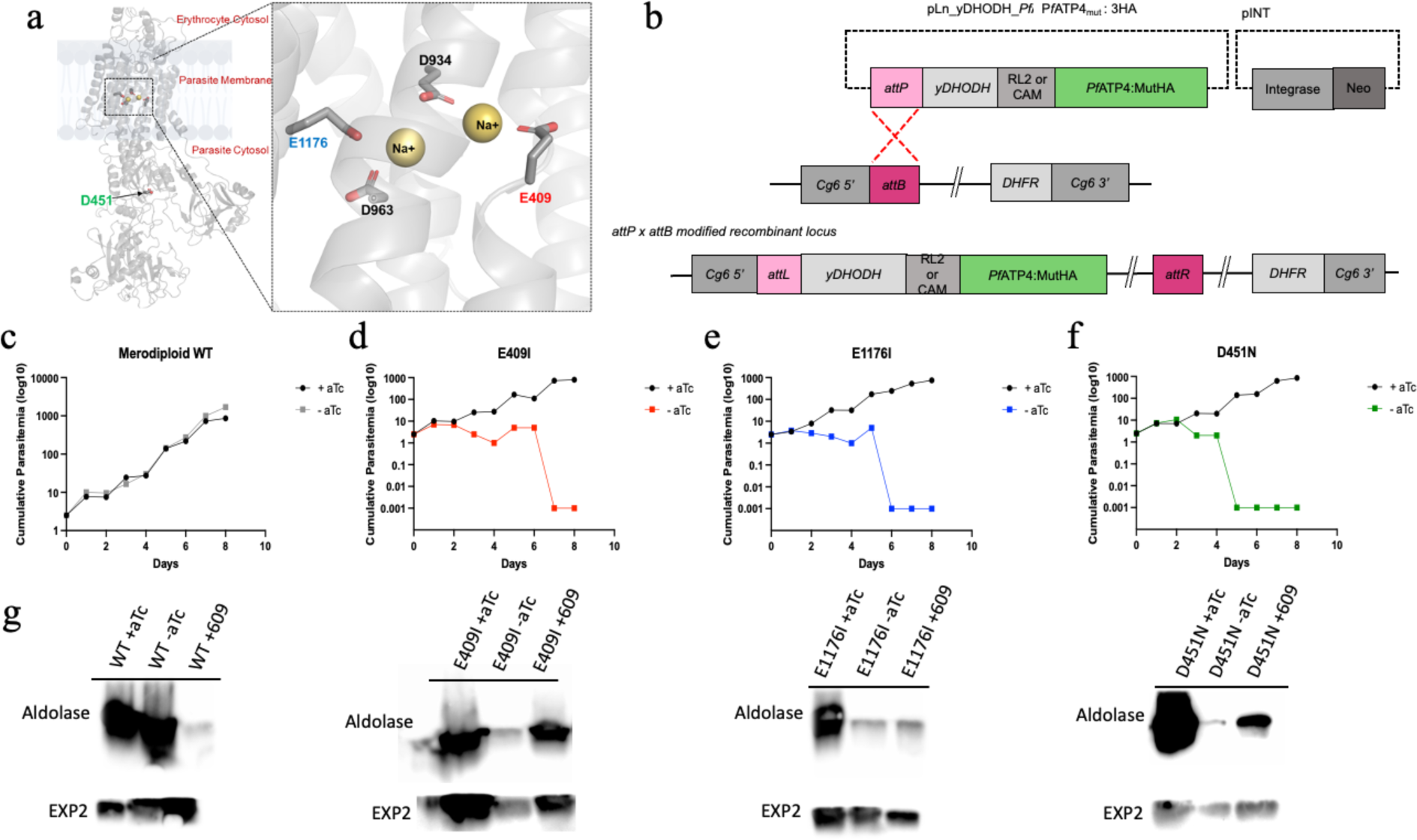
Assessing Predicted Na^+^ Coordination sites in P*f*ATP4. **(a)** Three-dimensional structure of the H_3_E2-ATP state of *Pf*ATP4 is depicted as ribbons and colored grey in a membrane environment with residues from Na^+^ coordination sites and phosphorylation site labeled and shown as licorice sticks and colored atom type (carbon: grey, nitrogen: blue and oxygen: red) with the inlet depicting the magnified Na^+^ binding site **(b)** Schematic representation of the integration strategy used that was used to generate merodiploid NF54*attB Pf*ATP4:3xMyc/*Pf*ATP4_(mut)_:3xHA parasites in which endogenous *Pf*ATP4 expression can be conditionally regulated by the TetR: DOZi aptamer system. The mutated *Pf*ATP4 allele is constitutively expressed under the control of the RL2 promoter from the cg6::attB locus. **(c-f)** Representative growth curves of (2 biological replicates) of merodiploid wild type, *Pf*ATP4_E409I_, *Pf*ATP4_E1176I_, and *Pf*ATP4_D451N_, parasites grown in the presence or absence of 250 nM aTc. Cultures were split 1:4 at 48, 96, and 144 hpi. Parasitemia was determined by counting at least 1000 red blood cells in each Giemsa-stained thin blood smear. Cumulative parasitemia was determined by multiplication of parasitemia and split factor (in this case, 4) over the time course. **(g)** Saponin sensitivity assay assessed by Western blot analysis of *Pf*ATP4_mut_ parasites grown with and without aTc for 60 hours, treated with 0.02% saponin and probed with an anti-aldolase antibody to measure cytosolic leakage. Parasites grown in aTc for 60 hours and then treated with 10 nM KAE609 prior to saponin treatment were used as a positive control.

The fact that these mutations could not support parasite growth suggests that there may be a defect in Na^+^ efflux by the mutated *Pf*ATP4s from the parasite cytosol. In a previous study, we demonstrated that the influx of Na^+^ due to *Pf*ATP4 inhibition led to the accumulation of cholesterol in the PPM (10). This effect is reflected by parasites acquiring sensitivity to treatment with saponin, a cholesterol-dependent detergent. The saponin sensitivity can be assessed by the leakage of cytosolic proteins, such as aldolase, after saponin treatment (10). We used this observation as a surrogate for assessing *Pf*ATP4 function. Parasites were grown in the presence and absence of aTc, and their sensitivity to saponin was assessed via western blot using an anti-aldolase antibody.

As a control, parasites were grown with aTc and treated with a *Pf*ATP4 inhibitor (KAE609) for 2 hours prior to saponin treatment. Merodiploid wild-type parasites (*Pf*ATP4_WT_) grown with or without aTc were not saponin sensitive, as indicated by the absence of aldolase leakage following saponin treatment **(Fig. 3g)**. In contrast, *Pf*ATP4_E409I_ and *Pf*ATP4_E1176I_ parasites grown without aTc showed evidence of cholesterol accumulation in the PPM, similar to parasites treated with KAE609 **(Fig. 3g)**. These results suggest that these mutations confer a functional defect to the protein.

To determine whether this defect could be due to dysregulation of Na^+^ coordination or improper protein localization to the PPM, we conducted an immunofluorescence assay (IFA) using an anti-HA antibody and an antibody to a known parasitophorous vacuolar membrane (PVM) marker, *Pf*EXP2 (30). The PPM, where *Pf*ATP4 localizes, is closely apposed to the PVM and, therefore, can be used to visualize proper protein localization. Similar to *Pf*ATP4_WT_, mutant proteins from both *Pf*ATP4_E409I_ and *Pf*ATP4_E1176I_ parasites were closely associated with *Pf*EXP2, indicating appropriate localization of the mutated *Pf*ATP4 **(Fig. 5)**. These data indicate that *Pf*ATP4_E409I_ and *Pf*ATP4_E1176I_ parasites confer a fitness defect likely due to dysregulation of Na^+^ homeostasis since the protein is properly localized to the PPM.

### Assessing a Potential Phosphorylation Site

P-type ATPases derive their name from an intermediate phosphorylated state that is shared by all proteins in this family (13, 14). Multiple X-ray crystal structures of intermediate states of the catalytic cycle of SERCA have revealed massive structural rearrangements of the cytoplasmic and TM domains(13, 18, 19) These structural rearrangements are determined by both the bound ion and the phosphorylation state of the pump (13). In P2-type ATPases, the phosphoacceptor during autophosphorylation is an aspartate side chain in a universally conserved DKTG motif in the P domain (13). To transition from the E1 unbound state to the 2Na^+^E1P-ADP state, ATP must be bound and hydrolyzed to ADP. For this, ATP is coordinated between the N and P domains, where the conserved aspartate breaks the interdomain linkage of ATP to create ADP (13).

Sequence alignments predict that residue D451 in *Pf*ATP4 is homologous to D351 in SERCA, which was identified to be part of the DKTG motif. To assess this, we created a merodiploid parasite line with a D451N mutation in *Pf*ATP4 to force the pump into a stable, dephosphorylated state. We then assessed the ability of *Pf*ATP4_D451N_ parasites to complement endogenous *Pf*ATP4 knockdown. In the absence of aTc, endogenous *Pf*ATP4 was not expressed, while ectopic expression was maintained **(Fig. S5d)**. However, *Pf*ATP4_D451N_ could not complement endogenous *Pf*ATP4 deficiency compared to the *Pf*ATP4_WT_ **(Fig. 3f)**. After evaluation of parasite growth, we next assessed *Pf*ATP4 function using the saponin sensitivity assay as described above. Western blot analysis demonstrated that *Pf*ATP4_D451N_ parasites grown without aTc showed evidence of cholesterol accumulation in the PPM similar to treatment with KAE609, indicating that the mutation conferred a functional defect to *Pf*ATP4 **(Fig. 3g)**. Next, we conducted IFA to localize the *Pf*ATP4_D451N_ mutant protein. In contrast to the Na^+^ coordination site mutations described above, *Pf*ATP4_D451N_ failed to localize to the PPM, as shown by IFA analysis **(Fig. 5)**, suggesting that D451 may be required for proper protein translocation in addition to being critical for phosphorylation/de-phosphorylation cycling.

### Assessing *Pf*ATP4 Inhibitor Resistance-Associated Mutations

Although *Pf*ATP4 is an attractive target for multiple chemical scaffolds, resistance to these compounds arises relatively easily *in vitro*. So far, over twenty different mutations conferring resistance to various *Pf*ATP4 inhibitors have been identified (1–3, 31). Most of these mutations do not seem to confer a fitness cost to the parasite. However, one exception is the P412T mutation, which confers resistance to spiroindolone and DHIQ classes of *Pf*ATP4 inhibitors (3). Previous studies generated P412T mutant parasites through continuous drug exposure. Thus, a possibility remains that the observed fitness defect may be due to a mutation elsewhere in the genome. To assess this possibility, we generated a merodiploid line bearing a P412T mutation expressed from the ectopic locus (*Pf*ATP4_P412T_). The ability of the mutated allele to complement endogenous *Pf*ATP4 knockdown was tested by the withdrawal of aTc **(Fig. 4a)**. While drug-selected resistant parasites bearing a P412T mutation were reported to remain alive, albeit with diminished growth [3], merodiploid parasites with P412T mutation ceased growth upon the removal of aTc. The reasons for this discrepancy are not clear but may indicate a compensatory adjustment in parasites that were selected through continuous drug selection. We also engineered a merodiploid line expressing *Pf*ATP4 with two mutations, one known to confer resistance to pyrazoleamide (V178I) and the other to spiroindolone (G223R). Withdrawal of aTc did not affect the parasite’s growth phenotype **(Fig. 4b)**. Immunofluorescence examination showed that in *Pf*ATP4_P412T_ and *Pf*ATP4_V178I+G223R_ parasites, *Pf*ATP4 localized to the parasite surface, indicating appropriate localization of the protein **(Fig. 5)**

**Figure 4:**
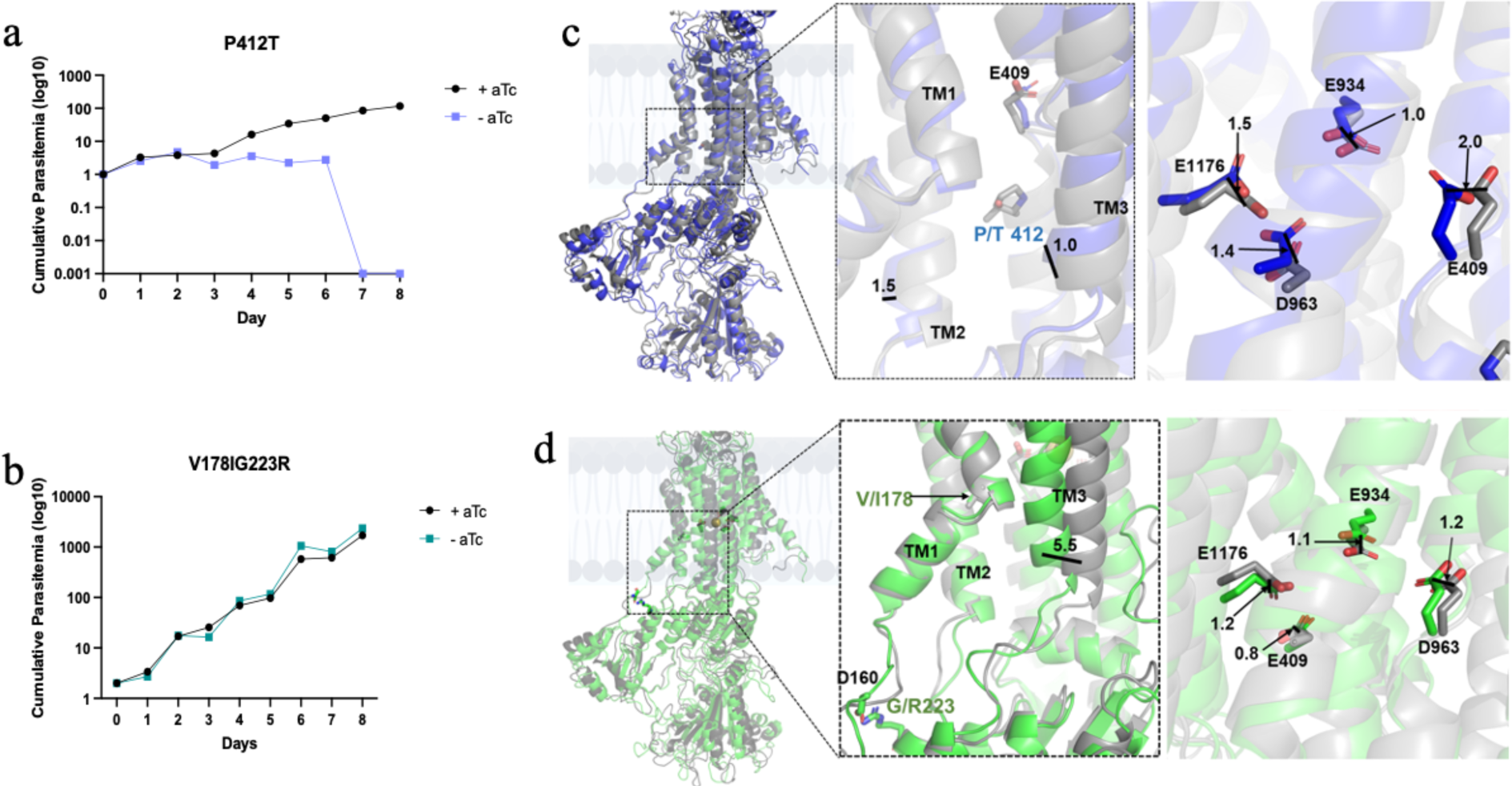
Structural Basis for Resistance-Associated Mutations of P*f*ATP4. **(a,b)** Representative growth curves of (2 biological replicates) of *Pf*ATP4_P412T_ and *Pf*ATP4_V178I+G223R_ parasites grown in the presence or absence of 250 nM aTc. Cultures were split 1:4 at 48, 96, and 144hpi. Parasitemia was determined by counting at least 1000 red blood cells in each Giemsastained thin blood smear. Cumulative parasitemia was determined by multiplication of parasitemia and split factor (in this case, 4) over the time course. **(c-d)** Structural superpositioning of wildtype H3E2-ATP of *Pf*ATP4 (represented as grey ribbons in all the models) and the P412T (represented as blue ribbons) (c) and V178I+G223R (represented as green ribbons) (d) mutant models are shown. In the magnified panels, the residues at the ions binding sites and the mutation sites are labeled and represented as licorice sticks and colored atom type with – nitrogen: blue, oxygen: red. The relative displacement of the helices or residues is indicated as black lines, and the displacement is measured in A.

To understand the structural consequences of P412T and V178I+G223R mutations, the H_3_E2-ATP structure from MD, which was previously simulated for 1μs to apparent equilibrium, was used. The homology models of the mutants were simulated for an additional 100 ns. A comparison of the RMSD plots suggested that the structures were stably equilibrated throughout the simulations **(Fig. S2a).** Superpositioning of these mutant models onto the wild-type H_3_E2-ATP structure showed RMSDs of 2.91 and 3.51 Å for the P412T and V178I+G223R models, respectively **(Table S2)**. Analysis of the mutant models showed significant movement of TM3 towards TM1 by 5.5 Å in the V178I+G223R mode **(Fig 4 c-d)**. This movement may lead to a decrease in the volume of the *Pf*ATP4 inhibitor binding cavity, which is lined by several residues from TM1 and TM3 regions, and may explain the loss of efficacy and drug resistance observed with these mutants. G223 lies in the cytoplasmic region of the *Pf*ATP4 structure, and the double mutation V178I+G223R results in the formation of a new salt bridge interaction between D160 and R223. This is a significant structural change that might impact the overall conformation of the E2 state, affecting drug sensitivity.

To understand the effects of these mutations on the Na^+^ coordination sites, the relative displacements of E409, E934, D963, and E1176 were measured **(Fig. 4 c-d).** A previous study on SERCA showed that the movement of E309 (corresponding to E409 in *Pf*ATP4) is critical for the E2 to E1 transition (22). In the P412T mutant, the displacement of E409 was ∼2Å, which was greater than the shift seen in the V178I+G223R mutant. The movement of E409 in the mutants could alter the H-bond network in the vicinity of E409, particularly with V404, N959, and N207, and affect the stability of the E2 state **(Fig. S2b)**. The H-bond distances between the protonated side chain of E409 and the V404 backbone were 2.2, 2.8, and 1.8 Å for the wild type, P412T, and V178I+G223R, respectively **(Fig. S2b-d)**. There was a significant change in the H-bond distances between the carbonyl backbone of E409 and N207 in the P412T mutant compared to other mutants. This H-bond exists only in the E2 state and is broken in the E1 state in the *Pf*ATP4 models **(Fig. S2c)**. Thus, based on these modeling results, we predict that the P412T mutation may decrease the stability of the E2 state and affect the E1/E2 equilibrium, potentially affecting parasite fitness.

#### Response of Merodiploid Parasites to Antimalarials

We used these merodiploid lines to assess their responses to three different antimalarial compounds (KAE609, PA21A092, and artemisinin) in the presence and absence of aTc. As shown in **Table 1**, Na^+^ coordination mutations (*Pf*ATP4_E409I_ and *Pf*ATP4_E1176I_), as well as phosphorylation cycle mutant (*Pf*ATP4_D451N_), showed hypersensitivity to treatment with *Pf*ATP4 inhibitors in the absence of aTc. Their susceptibility to artemisinin, however, was the same in the presence and absence of aTc. These results reveal synthetic lethality between diminished *Pf*ATP4 function and its inhibitors.

**Table 1:** Drug susceptibility of parental and merodiploid line(s) to *Pf*ATP4 inhibitors (KAE609 and PA21A092) and artemisinin were quantified using the hypoxanthine incorporation assay.

| Mutation | Condition | Compound (EC50) |  |  |
| --- | --- | --- | --- | --- |
|  |  | KAE609<br>(spiroindolone) | PA21A092<br>(Pyrazoleamide) | Artemisinin |
| **Parental Line**<br>(NF45attB<br>*Pf*ATP4:3xMyc) | + aTc | 0.313 ± 0.092 | 5.69 ± 0.61 | n/a |
| **Merodiploid**<br>**WT** | + aTc | 0.605 ± 0.034 | 8.6 ± 0.42 | 14.30 ± 0.4 |
|  | - aTc | 0.245 ± 0.021 | 3.09 ± 0.35 | 14.0 ± 0.57 |
| **E409I**<br>(Na<sup>+</sup> Coordination) | + aTc | 0.181 ± 0.018 | 2.97 ± 0.32 | 12.79 ± 0.8 |
|  | - aTc | 0.026 ± 0.024 | 0.53 ± 0.38 | 14.88 ± 1.7 |
| **E1176I**<br>(Na<sup>+</sup> Coordination) | + aTc | 0.246 ± 0.042 | 5.3 ± 0.96 | 13.71 ± 0.73 |
|  | - aTc | 0.049 ± 0.020 | 0.82 ± 0.05 | 13.53 ± 0.44 |
| **D451N**<br>(Phosphorylation<br>Cycle) | + aTc | 0.077 ± 0.004 | 3.62 ± 0.36 | 14.69 ± 0.56 |
|  | - aTc | 0.008 ± 0.011 | 1.16 ± 0.44 | 14.06 ± 0.09 |
| **P412T**<br>(Resistance to<br>Spiroindolones and<br>DHIQ's) | +aTc | 0.405 ± 0.031 | 8.6 ± 0.21 | 13.46 ± 1.7 |
|  | - aTc | 0.061 ± 0.005 | 1.56 ± 0.21 | 24.79 ± 1.4 |
| **V178IG223R**<br>(Resistance to<br>Pyrazoleamides and<br>Spiroindolones) | + aTc | 2.38 ± 0.053 | 54.23 ± 8.3 | 12.9 ± 1.1 |
|  | - aTc | 4.28 ± 0.18 | 135.8 ± 5.6 | 11.3 ± 4.0 |

Resistance-associated mutations were also assessed for their response to drug treatment. *Pf*ATP4_V178I+G223R_ parasites were resistant to both KAE609 and PA21A092, as expected. However, *Pf*ATP4_P412T_ parasites, which express the P412T mutation, showed *hypersensitivity* to both KAE609 and PA21092. This discrepancy could be due to a compensatory adjustment in drug pressure-selected parasites bearing a P412T mutation, which would not be present in our merodiploid line.

## Discussion

Malaria parasites encode thousands of proteins, a large percentage of which are essential for parasite survival (32, 33). In the search for new antimalarials to counter the ever-present threat of resistance emergence, structural and functional details of potential targets are of great value. However, at present, such details are available only for a handful of proteins encoded by malaria parasites. Here, we built a robust homology model for a validated antimalarial drug target *Pf*ATP4 using extensive MD simulations based on four principal catalytic states of the Ca^2+^ pump, SERCA **(Fig.1b-d)**. This dynamic model predicted motions of various domains of *Pf*ATP4 as it undergoes the ion pumping cycle, as well as critical residues involved in its function. While the AI-based tool AlphaFold is remarkable in providing models of proteins for which homologous structures are not available, the models it predicts are static and do not take into consideration the explicit environment in which the proteins exist. Therefore, the approach we used here has a significant advantage in gaining structural information on proteins for which authentic structures are not available. Clearly, since these are models, their predictions would need to be experimentally validated.

Such experimental validation for *Pf*ATP4, however, poses challenges since expression of this protein in its functionally active form in heterologous system has not been possible thus far. Since *Pf*ATP4 is an essential gene in *P. falciparum*, its mutational analysis could be impeded due to the haploid nature of the parasite genome. To circumvent this, we developed a system to assess the phenotypic consequences of site-directed mutagenesis of *Pf*ATP4 by generating merodiploid parasites in which two alleles of *Pf*ATP4 are present **(Fig. 2e)**. The endogenous allele on chromosome 12 was engineered to be conditionally expressed using the TetR-DOZI/aptamer regulation, whereas the second allele was constitutively expressed from an ectopic locus on chromosome 7 of *P. falciparum.* The endogenous allele was tagged with a c-Myc epitope, and the second allele was tagged with a 3x-HA epitope. In addition, we used the NF54 parasites for generating transgenic lines with ectopic *Pf*ATP4 allele from the Dd2 parasites that had a non-synonymous polymorphism, which permitted distinction between the encoded proteins when subjected to immunoprecipitation using epitope-specific antibodies **(Fig. 2i-j)**. Lack of co-immunoprecipitation, as well as proteomic analyses, showed that *Pf*ATP4 molecules encoded by the two alleles did not interact with each other **(Fig. 2h)**. Thus, the function of the complementing *Pf*ATP4 allele could be independently evaluated when expression of the endogenous allele was suppressed by withdrawing aTc from the culture medium.

Our model predicted acidic residues within *Pf*ATP4 that coordinate Na^+^ as it is being pumped across the parasite plasma membrane. We tested these predictions in two merodiploid lines, each expressing mutated *Pf*ATP4 bearing single amino acid change involved in Na^+^ coordination (E409I and E1176I). Unlike the wild-type *Pf*ATP4 allele, these mutant alleles failed to complement the knockdown of endogenous *Pf*ATP4 **(Fig. 3c-e)**. Immunofluorescence analysis showed the mutated protein localized to the parasite surface, indicating that there was no apparent defect in the transport of the protein **(Fig. 5)**. We also examined the mutation of the predicted conserved aspartate (D451N) involved in the phosphorylation cycle of the pump. These parasites exhibited a relatively more severe growth inhibition phenotype when expressing this allele in the absence of the endogenous allele **(Fig. 3f)**. Interestingly, unlike the mutations involving Na^+^ coordination residues, this mutation resulted in the mislocalization of *Pf*ATP4 in the cytoplasm **(Fig. 5)**, suggesting a possible link between the phosphorylation state of the protein and its appropriate transport to the parasite surface.

**Figure 5:**
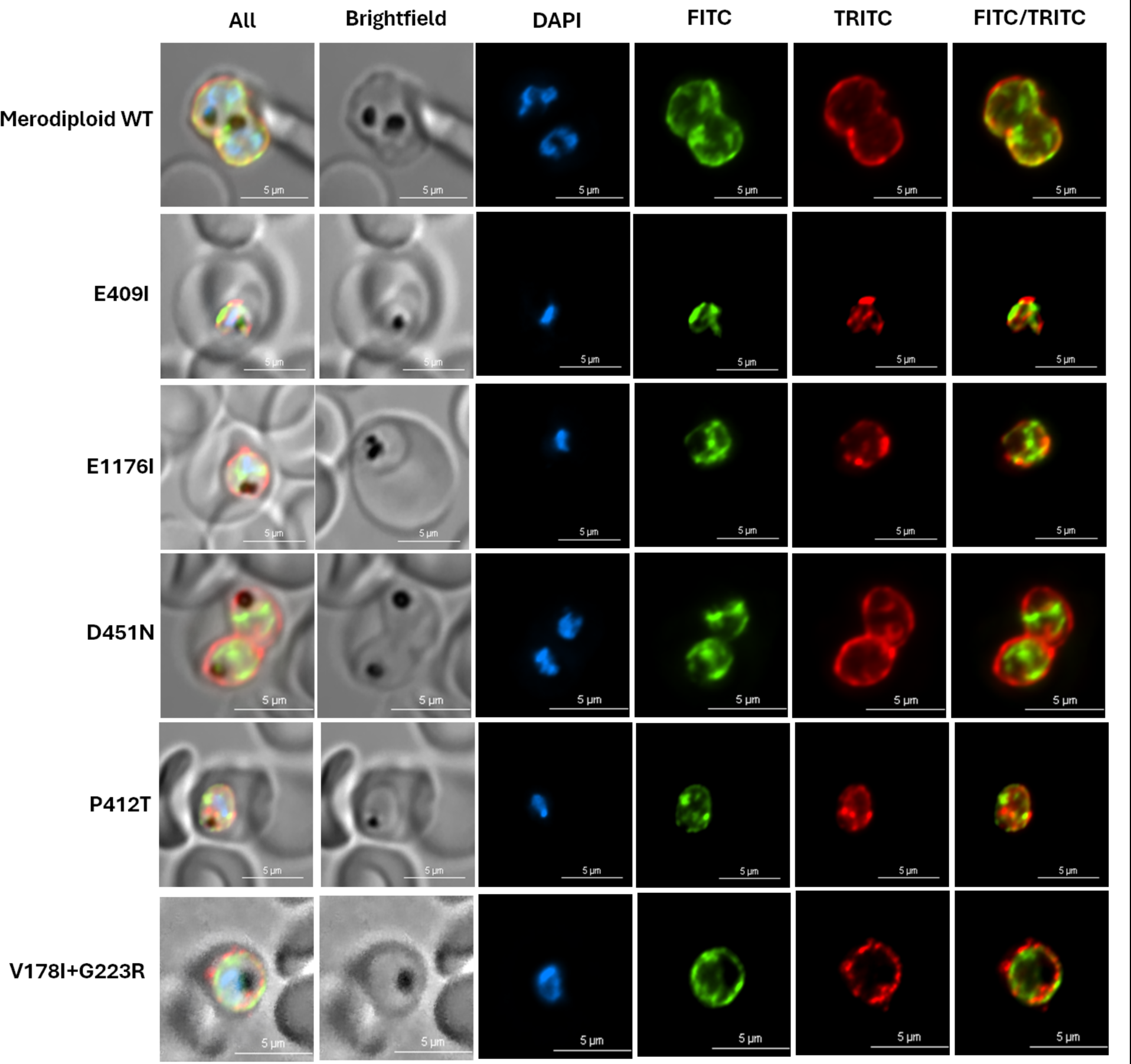
Immunofluorescence assays were conducted to observe the localization of ectopic *Pf*ATP4_Mut_:3xHA from (all lines). Parasites were fixed and stained with DAPI (blue), anti-HA antibody (green), and anti-EXP2 antibody (red). Images are representative of ∼20 captured images. Scale bars denote 5um.

In a previous study, we have shown that inhibition of *Pf*ATP4 for a short 2 h period results in massive changes in parasite morphology and lipid homeostasis (10). One such change involves reversible accumulation of cholesterol in the parasite plasma membrane, which can be assessed as leakage of cytosolic proteins following treatment with cholesterol-dependent detergents such as saponin (10). We have also found that inhibition of another pump, *Pf*NCR1 (*P. falciparum* Niemann-Pick type C1-related), also results in a similar accumulation of cholesterol in the parasite plasma membrane observed as saponin sensitivity (34). Since *Pf*NCR1 has a sterol-binding domain, it is predicted to be involved in cholesterol homeostasis in the parasite and, thus, likely to be secondarily inhibited following *Pf*ATP4 inhibition (35). As shown in **Fig. 3g**, mutations that are predicted to affect *Pf*ATP4 function also result in the induction of saponin sensitivity in merodiploid parasites. We hypothesize that *Pf*NCR1 requires a Na^+^ gradient across the parasite plasma membrane to maintain cholesterol homeostasis and that a collapsed Na^+^ gradient in *Pf*ATP4 mutant parasites leads to inhibition of *Pf*NCR1, resulting in cholesterol accumulation in the parasite plasma membrane.

The utility of the homology model we have built was tested to assess the structural consequences of drug resistance-associated mutations in *Pf*ATP4. We engineered two merodiploid lines expressing such mutations and assessed their growth drug resistance phenotypes. Parasites bearing V178I+G223R mutations did not show changes in their growth phenotype and showed increased resistance to drugs when grown in the absence of aTc. Parasites bearing the P412T mutation, however, showed a severe growth defect when grown in the absence of aTc and were hypersensitive to *Pf*ATP4 inhibitors. MD simulations of this mutation revealed an impact on the relative stability of the intermediate states, which could explain the phenotype observed.

In summary, we have combined MD simulations and genetic manipulations of an antimalarial drug target to gain insights into its function. This approach also provides insights into the structure-activity relationship for hundreds of compounds synthesized as part of an effort to develop antimalarials that target *Pf*ATP4, which will be described elsewhere (Nepal et al.)

## MATERIALS AND METHODS

### Homology Modeling

The amino acid sequence of *Pf*ATP4 was obtained from the Uniprot database (36) (accession number: Q9U445) and is comprised of 1264 residues. A position-specific BLAST suggested the human SERCA protein as the closest structural homolog. Since SERCA undergoes significant conformational changes across its pumping cycle (**Fig. 1b**), we modeled the analogous conformational states of the *Pf*ATP4. The structure of each *PfATP4* conformational state was generated using the homology modeling module of Molecular Operating Environment software (MOE, ver 10.1 (37)) using the corresponding crystal structures of the SERCA as templates. The 2Na^+^E1P-ADP was modeled with the SERCA structure (PDB:3BA6 (38)) with the Na^+^ ion coordinated by E409, E934, D963, and E1176, which were modeled in the deprotonated state. In this state, D451 is phosphorylated, and one ADP molecule is bound in the nucleotide-binding domain. ADP molecule was mimicked by AMP phosphoramidate as in the SERCA crystal structure. The E2P conformational state was modeled with SERCA E2P structure (PDB:3B9B) as a template with D451 in the phosphorylated state but without any Na^+^ ions. The H_3_E2P-ATP conformational state was modeled with the corresponding SERCA structure (PDB:3FGO) with D451 as phosphorylated and with one ATP molecule bound to the nucleotide-binding domain and E409, E934, and E1176 in their protonated states. The H_3_E2-ATP conformation was modeled with the corresponding SERCA structure (PDB:2C88 (40)) as a template. In this state, D451 was dephosphorylated and bound to the ATP binding domain wherein ATP was modeled as the phosphomethylphosphonic acid adenylate ester (ACP) and E409, E934 and E1176 were protonated.

### All-atom MD simulations

The models were equilibrated with all-atom MD simulations in a membrane environment using NAMD software (41). A membrane patch composed of 70% 1-palmitoyl-2-oleoyl-sn-glycero-3-phosphocholine (POPC), 20% 1-palmitoyl-2-oleoylphosphatidylethanolamine (POPE), 5% cholesterol and 5% Phosphatidylinositol 4,5-bisphosphate (PIP2) was simulated. The number of lipids and the corresponding lipid patch dimension for each system are detailed in **Table S2.** The system was modeled with an NPT ensemble using the Nose-Hoover thermostat and the MTK barostat, and the protein, lipid, and other heteroatoms representing ATP and ADP were represented using the CHARMM36 Force field (42)and TIP3P model for water, and the entire system was neutralized with 0.15M KCl. MD simulations for each system were carried out for a production run of 115ns with 2fs time step. The H_3_E2-ATP structure was simulated for an additional 1µs on the Anton2 supercomputer (43) with the same force field and time step. The force-field parameters for AMP phosphoramidate and phosphomethylphosphonic acid adenylate ester (ACP), which mimics ADP and ATP molecules, were generated from the CGennFF program (44). P412T and G223R+V178I mutants were constructed from the H_3_E2-ATP state using the final conformation of the 1µs simulation. The mutated systems were then simulated for an additional 100 ns on the Anton2 supercomputer (43). In the wild-type and mutant model simulations, the ACP molecule, which mimics ATP, was constrained in its position with a harmonic restraining potential of 5kcal/mol, which was released in the final steps of the simulation.

### *P. falciparum* culture maintenance

Asexual P. falciparum parasites were cultured in commercially obtained Human O+ erythrocytes and maintained at 2.5% hematocrit in RPMI 1640 medium supplemented with 0.5% w/v Albumax II (Gibco by ThermoFisher Scientific), sodium bicarbonate (2.1g/liter, Corning by ThermoFisher Scientific), HEPES (15mM, MilliporeSigma), hypoxanthine (10mg/liter, Fisher Scientific), gentamycin (50mg/liter, VWR), and further supplemented with selection drugs as needed. Parasite cultures were maintained at ∼5% parasitemia or lower (unless otherwise specified) under an atmosphere of 90% N_2_, 5% O_2_, and 5% CO_2_. Cultures were maintained at 5% parasitemia or lower unless otherwise specified. Periodically, mycoplasma testing was done to ensure that cultures were mycoplasma negative.

### Generation of NF54*attB Pf*ATP4:3xMyc Transgenic Line

To generate conditional knockdown of endogenous NF54 *Pf*ATP4:3xMyc tagged parasites, we utilized the TetR-Dozi system. The original pMG75-ATP4 vector (29) was a gift from Jacquin Niles (MIT). To allow appropriate single cross-over recombination at the endogenous locus and to facilitate the development of merodiploid constructs, the original vector was modified to remove the *attP* site (45). This plasmid was used to transfect NF54*attB* parasites as previously described (46). The transfected parasites were cloned by limited dilution in the presence of aTc (250nM) and in the absence of the selection drug, blasticidin S (BS-2.5.g/ml).

### Generation of Merodiploid Parasite Lines

To generate merodiploid parasite lines, full-length *Pf*ATP4 was PCR amplified from Dd2 wild-type parasite genomic DNA and cloned into pSC-B, a PCR blunt-end cloning vector, using the StrataClone Blunt PCR cloning kit (Agilent Technologies). After Sanger sequencing verification of selected clones, various mutations of *Pf*ATP4 were made using the oligonucleotides, as shown in **Supplementary Table S4**. Following sequence confirmation of the mutations, the pSC-B *Pf*ATP4_mut_ was digested (AvrII+BsiWI) and ligated into a pLN-yDHOD *attP* vector (modified from pLN-ENR-GFP) (47) digested at the same sites and having either a calmodulin or RL2 promoter to drive expression of *Pf*ATP4 **(Fig. S3)**. The final transfection vectors were verified by whole plasmid nanopore sequencing. Merodiploid parasite lines were generated by transfection of NF54*attB Pf*ATP4:3xMyc parasites with pLN vectors containing mutations of *Pf*ATP4 along with an integrase plasmid. Transfection vectors were co-precipitated with the integrase plasmid (47) using 0.5M sodium acetate (pH 7) and ethanol. Transfections were carried out as previously described (45). After electroporation, parasites were cultured in media supplemented with 250nM aTc for 48 hours, at which time, G418 (10 .g/ml), DSM1(1.5 .g/ml), and BS (2.5 .g/ml, InvivoGen) were added. After one week, G418 was removed, and parasites were routinely maintained in media supplemented with aTc, BS, and DSM1. Proper integration at the *attB* site was assessed using the primers listed in **Supplementary Table S4** (47) and depicted in **Fig. S4**.

The generation of the *Pf*ATP4_V178I+G223R_ parasite line differed slightly from the other merodiploid lines. Genomic DNA from Cmpd 2-1 resistant parasites was PCR amplified to obtain the *Pf*ATP4_V178I_ mutant gene (2). This DNA was then ligated into a PSC vector at the AvrII+BsiWI sites. Following this, primers listed in Supplemental Table S4 were used to engineer a G223R mutation in *Pf*ATP4, resulting in a V178I+G223R double mutant parasite line (*Pf*ATP4_V178I+G223R_).

### Parasite Proliferation Assays

To assess for phenotypic defects due to *Pf*ATP4 mutations, transgenic parasites were synchronized by osmotic lysis of late-stage parasites using 0.5M alanine. Ring-stage parasites were washed twice in RPMI 1640 medium and centrifuged at 1500xg for 10 minutes to remove residual aTc. Parasites were then split into two flasks, each containing a starting parasitemia of 1%, and grown in media with or without aTc for eight days. Cultures were split at the trophozoite stage, with 75% of the culture collected for Western Blot analysis and 25% retained to continue parasite growth. Cumulative parasitemia was calculated from day 0 to day 8 and plotted using GraphPad Prism.

### *Pf*ATP4 antibody generation

*Pf*ATP4 sequences encoding amino acids 449-760 were amplified by PCR and cloned into pET28a to generate His-tagged protein in *E. coli* BL21Codonplus (DE3) RIL bacteria (Agilent Technologies) bacteria. The protein was isolated using a Ni-NTA column (Qiagen). 25 μg of isolated protein was used to immunize mice by subcutaneous injection with complete Freund’s adjuvant and boosted three times with incomplete Freund’s adjuvant to generate *Pf*ATP4 antibody.

### Western Blot Analysis

Trophozoite stage parasites were lysed using 0.05% Saponin in 1X PBS supplemented with 1μg/mL protease inhibitor cocktail (P8215, MilliporeSigma). Parasites were spun down at 1500 xg for 10 minutes at room temperature. Parasite pellets were washed twice in 1X PBS until the lysate was clear. After washing, the pellets were resuspended in 100 µL of a solution containing 1% sodium dodecyl sulfate (SDS), 5% β-mercaptoethanol, and 1% bromophenol blue, mixed by vigorous pipetting and stored at -20°C. The lysate was spun down at 17000 xg for 10 minutes, and the resulting supernatant was used for SDS-polyacrylamide gel electrophoresis (SDS-PAGE). 40μL of each sample was loaded into each lane, and electrophoresis was performed using a standard protocol. Membranes were blocked with 5% w/v fat-free milk powder for 90 minutes.

Blots were then incubated with monoclonal anti-mouse HA antibody (F-7, sc-7392, Santa Cruz Biotechnology, Lot # 12923) at 1:10,000 dilution or monoclonal anti-mouse Myc antibody (9B11, Cell Signaling 2276S) at 1:10,000 followed by incubation with either secondary goat pAb to mouse IgG HRP (ab97040, Abcam, Lot # GR3219575-4) at 1:10,000 or secondary mouse antibody to rabbit IgG HRP (Santa Cruz Biotechnology, Lot # L1819). As a loading control, blots were later probed with a rabbit anti-PyEXP2 primary antibody (a gift from Dr. James Burns, Drexel University) at 1:10,000, followed by a secondary mouse antibody to rabbit IgG HRP (see previous) at 1:10,000. Blots were then imaged on a Bio-Rad ChemiDoc™MP Imaging System.

### Saponin Sensitivity Assay

Ring-stage parasites were tightly synchronized with 0.5M alanine and split into three T-25 flasks; aTc was removed from one of these. After 60 h, one flask grown in the presence of aTc was treated with 10 nM of KAE609 for 2 h at 37°C. To monitor sensitivity to saponin, trophozoite stage parasites were released from erythrocytes using 0.02% Saponin in 1X PBS supplemented with 1 μg/mL protease inhibitor cocktail (P8215, MilliporeSigma). Parasites were centrifuged at 1500xg for 10 minutes at room temperature. Parasite pellets were washed three times in 1X PBS until the lysate was clear. After washing, the pellet was resuspended in ∼ five volumes of RIPA buffer supplemented with 1µg/ml of protease inhibitor cocktail (P8215, MilliporeSigma). The samples were further analyzed via western blot for leakage of cytosolic aldolase. Blots were incubated with anti-rabbit *Plasmodium* aldolase HRP antibody (ab38905, Abcam, Lot # GR289448-4) at 1:10000 dilution and processed as described above.

### Immunofluorescence Assays

Trophozoite stage parasites were fixed in 4% paraformaldehyde + 0.001% glutaraldehyde and rotated at 37°C for 1 h and then 4°C overnight. Samples were washed with 1X PBS + 0.001% Tween and spun down at 3000 rpm for 5 minutes. Samples were then permeabilized with 0.25% Triton X-100 and rotated at room temperature for 10 minutes. Following permeabilization and washing, samples resuspended at ∼100 mg/mL were reduced with 0.1 mg/mL NaBH_4_ while rotating at room temperature for 5 minutes. Following washing, samples were blocked w/v 3% BSA for 90 minutes and then incubated with monoclonal anti-mouse HA antibody (F-7, sc-7392, Santa Cruz Biotechnology, Lot # 12923) at 1:500 and rabbit anti-P*y*EXP2 primary antibody (a gift from Dr. James Burns, Drexel University) at 1:500 followed by goat anti-mouse IgG H+L Alexa Fluor™Plus 488 (Invitrogen, A32723, Lot # SG251135) at 1:500 and goat anti-rabbit IgG H+L AlexaFluor™ 568 (Invitrogen, A11036, Lot # 2447870) at 1:500. Samples were then equilibrated and resuspended in Gold Anti-Fade + DAPI (S2828, ThermoFisher Scientific) and mounted onto slides for microscopy. Parasites were then visualized, and z-stack images were captured using a Nikon Ti microscope. Images were then 2D deconvoluted and processed using Nikon NIS Elements (5.30.02) Imaging Software.

### Phylogenetic analysis

Sequences of P2D-type ATPases from species representing a broad range of eukaryotic groups were identified from previous studies (15, 48) and BLAST (49) searches of NCBI (50) and EupathDB (51) databases. The sequences chosen for alignment are listed in **Supplementary Table S1**. Details of the software packages and methodology used for phylogenetic analysis are given in *Supplemental Methods*.

### Growth inhibition assay

All parasite growth inhibition assays were performed in triplicate in 96-well plates as described by Desjardin *et al*. (52). *P. falciparum*-infected erythrocytes at 1.0% initial parasitemia and 1.5% hematocrit were exposed to various concentrations of the indicated drug/inhibitor 24 h and then pulsed with 0.5 μCi of ^3^H-hypoxanthine for 24 h. Untreated parasitized and unparasitized red blood cells were incubated concurrently with the treated parasites as controls for growth and radioactive precursor incorporation. The 96 well plates were then frozen at -80°C for 24 h to promote erythrocyte and parasite lysis. Following freezing, the plates were warmed to 37°C, and parasites were harvested onto EasyTab^TM^-C Self-Aligning Glass Fiber Filters (Packard, Meridian, CT, USA). The filters were then dried completely and placed into an Omnifilter^TM^ 96-well plate filter case (Packard, Meridian, CT, USA). In order to measure the beta-radiation of incorporated ^3^H-hypoxanthine, 30μl of OmniScint^TM^ (Packard, Meridian, CT, USA) scintillation fluid was added to each well of the Omnifilter case. The Omnifilter was then covered with a TopSeal and counted with a TopCount^TM^ radiation counter. The incorporation of radioactivity into nucleic acids served as a measure of cell proliferation and is calculated by TopCount as counts per minute (cpm). The total cpm of the non-parasitized red blood cells was subtracted from the cpm of all other wells. The percent growth was plotted using GraphPad Prism software by graphing the cpm versus the drug/inhibitor concentration. The cpm of each sample was converted to percent by dividing it by the cpm of the untreated control. The best-fit curve was calculated by adjusting the lowest percent to equal zero and the highest percent to 100. From this best-fit curve, the concentration of the inhibitor that inhibits 50% of the parasite growth (IC_50_) is calculated.

## Acknowledgements

This work was supported by grants from National Institutes of Health R01AI154499 (to ABV and SK) and R01AI132508 (to ABV), and Anton 2 award MCB210021P (to SK). Anton 2 computer time was provided by the Pittsburgh Supercomputing Center (PSC) through Grant R01GM116961 from the National Institutes of Health. The Anton 2 machine at PSC was generously made available by D.E. Shaw Research.

## References

1. M. Rottmann et al., Spiroindolones, a potent compound class for the treatment of malaria. Science 329, 1175–1180 (2010).

2. A. B. Vaidya et al., Pyrazoleamide compounds are potent antimalarials that target Na+ homeostasis in intraerythrocytic Plasmodium falciparum. Nat Commun 5, 5521 (2014).

3. M. B. Jimenez-Diaz et al., (+)-SJ733, a clinical candidate for malaria that acts through ATP4 to induce rapid host-mediated clearance of Plasmodium. Proc Natl Acad Sci U S A 111, E5455–5462 (2014).

4. N. J. Spillman et al., Na(+) regulation in the malaria parasite Plasmodium falciparum involves the cation ATPase PfATP4 and is a target of the spiroindolone antimalarials. Cell Host Microbe 13, 227–237 (2013).

5. A. S. M. Dennis, J. E. O. Rosling, A. M. Lehane, K. Kirk, Diverse antimalarials from whole-cell phenotypic screens disrupt malaria parasite ion and volume homeostasis. Sci Rep 8, 8795 (2018).

6. A. M. Lehane, M. C. Ridgway, E. Baker, K. Kirk, Diverse chemotypes disrupt ion homeostasis in the Malaria parasite. Mol Microbiol 94, 327–339 (2014).

7. N. J. White et al., Spiroindolone KAE609 for falciparum and vivax malaria. N Engl J Med 371, 403–410 (2014).

8. A. H. Gaur et al., Safety, tolerability, pharmacokinetics, and antimalarial efficacy of a novel Plasmodium falciparum ATP4 inhibitor SJ733: a first-in-human and induced blood-stage malaria phase 1a/b trial. Lancet Infect Dis 20, 964–975 (2020).

9. A. S. M. Dennis, A. M. Lehane, M. C. Ridgway, J. P. Holleran, K. Kirk, Cell Swelling Induced by the Antimalarial KAE609 (Cipargamin) and Other PfATP4-Associated Antimalarials. Antimicrob Agents Chemother 62 (2018).

10. S. Das et al., Na+ Influx Induced by New Antimalarials Causes Rapid Alterations in the Cholesterol Content and Morphology of Plasmodium falciparum. PLoS Pathog 12, e1005647 (2016).

11. H.-J. Apell, How do P-type ATPases transport ions? Bioelectrochemistry 63, 149–156 (2004).

12. K. B. Axelsen, M. G. Palmgren, Evolution of substrate specificities in the P-type ATPase superfamily. J Mol Evol 46, 84–101 (1998).

13. M. Dyla, M. Kjaergaard, H. Poulsen, P. Nissen, Structure and Mechanism of P-Type ATPase Ion Pumps. Annu Rev Biochem 89, 583–603 (2020).

14. M. G. Palmgren, P. Nissen, P-type ATPases. Annu Rev Biophys 40, 243–266 (2011).

15. B. Benito, B. Garciadeblas, P. Schreier, A. Rodriguez-Navarro, Novel p-type ATPases mediate high-affinity potassium or sodium uptake in fungi. Eukaryot Cell 3, 359–368 (2004).

16. A. Rodriguez-Navarro, B. Benito, Sodium or potassium efflux ATPase a fungal, bryophyte, and protozoal ATPase. Biochim Biophys Acta 1798, 1841–1853 (2010).

17. A. Fraile-Escanciano, B. Garciadeblas, A. Rodriguez-Navarro, B. Benito, Role of ENA ATPase in Na(+) efflux at high pH in bryophytes. Plant Mol Biol 71, 599–608 (2009).

18. C. Toyoshima et al., Crystal structures of the calcium pump and sarcolipin in the Mg2+-bound E1 state. Nature 495, 260–264 (2013).

19. W. Kuhlbrandt, Biology, structure and mechanism of P-type ATPases. Nat Rev Mol Cell Biol 5, 282–295 (2004).

20. C. Toyoshima, G. Inesi, Structural basis of ion pumping by Ca2+-ATPase of the sarcoplasmic reticulum. Annu Rev Biochem 73, 269–292 (2004).

21. C. Toyoshima, M. Nakasako, H. Nomura, H. Ogawa, Crystal structure of the calcium pump of sarcoplasmic reticulum at 2.6 A resolution. Nature 405, 647–655 (2000).

22. N. Tsunekawa, H. Ogawa, J. Tsueda, T. Akiba, C. Toyoshima, Mechanism of the E2 to E1 transition in Ca^2+^ pump revealed by crystal structures of gating residue mutants. Proceedings of the National Academy of Sciences 115, 12722–12727 (2018).

23. C. Toyoshima, T. Mizutani, Crystal structure of the calcium pump with a bound ATP analogue. Nature 430, 529–535 (2004).

24. C. Olesen et al., The structural basis of calcium transport by the calcium pump. Nature 450, 1036–1042 (2007).

25. A. Nagarajan, J. P. Andersen, T. B. Woolf, Coarse-Grained Simulations of Transitions in the E2-to-E1 Conformations for Ca ATPase (SERCA) Show Entropy–Enthalpy Compensation. Journal of Molecular Biology 422, 575–593 (2012).

26. R. Aguayo-Ortiz, L. M. Espinoza-Fonseca, Linking Biochemical and Structural States of SERCA: Achievements, Challenges, and New Opportunities. International Journal of Molecular Sciences 21, 4146 (2020).

27. W. Tian, C. Chen, X. Lei, J. Zhao, J. Liang, CASTp 3.0: computed atlas of surface topography of proteins. Nucleic Acids Research 46, W363–W367 (2018).

28. J. Jumper et al., Highly accurate protein structure prediction with AlphaFold. Nature 596, 583–589 (2021).

29. S. M. Ganesan, A. Falla, S. J. Goldfless, A. S. Nasamu, J. C. Niles, Synthetic RNA-protein modules integrated with native translation mechanisms to control gene expression in malaria parasites. Nat Commun 7, 10727 (2016).

30. E. Meibalan et al., Host erythrocyte environment influences the localization of exported protein 2, an essential component of the Plasmodium translocon. Eukaryot Cell 14, 371–384 (2015).

31. A. H. Lee, D. A. Fidock, Evidence of a Mild Mutator Phenotype in Cambodian Plasmodium falciparum Malaria Parasites. PLoS One 11, e0154166 (2016).

32. M. Zhang et al., Uncovering the essential genes of the human malaria parasite Plasmodium falciparum by saturation mutagenesis. Science 360 (2018).

33. E. Bushell et al., Functional Profiling of a Plasmodium Genome Reveals an Abundance of Essential Genes. Cell 170, 260–272 e268 (2017).

34. S. Bhatnagar, S. Nicklas, J. M. Morrisey, D. E. Goldberg, A. B. Vaidya, Diverse Chemical Compounds Target Plasmodium falciparum Plasma Membrane Lipid Homeostasis. ACS Infect Dis 5, 550–558 (2019).

35. E. S. Istvan et al., Plasmodium Niemann-Pick type C1-related protein is a druggable target required for parasite membrane homeostasis. Elife 8 (2019).

36. T. U. Consortium, UniProt: the Universal Protein Knowledgebase in 2023. Nucleic Acids Research 51, D523–D531 (2022).

37. Anonymous, Molecular Operating Environment (MOE), 2022.02 Chemical Computing Group ULC, 1010 Sherbooke St. West, Suite #910, Montreal, QC, Canada, H3A 2R7, 2023.

38. C. Olesen et al., The structural basis of calcium transport by the calcium pump. Nature 450, 1036–1042 (2007).

39. M. Laursen et al., Cyclopiazonic Acid Is Complexed to a Divalent Metal Ion When Bound to the Sarcoplasmic Reticulum Ca2+-ATPase*. Journal of Biological Chemistry 284, 13513–13518 (2009).

40. A.-M. L. Jensen, T. L.-M. Sørensen, C. Olesen, J. V. Møller, P. Nissen, Modulatory and catalytic modes of ATP binding by the calcium pump. The EMBO Journal 25, 2305–2314 (2006).

41. L. V. Kalé, A. Bhatele, E. J. Bohm, J. C. Phillips, "NAMD (NAnoscale Molecular Dynamics)" in Encyclopedia of Parallel Computing, D. Padua, Ed. (Springer US, Boston, MA, 2011), 10.1007/978-0-387-09766-4_505, pp. 1249–1254.

42. R. B. Best et al., Optimization of the additive CHARMM all-atom protein force field targeting improved sampling of the backbone phi, psi and side-chain chi(1) and chi(2) dihedral angles. J Chem Theory Comput 8, 3257–3273 (2012).

43. D. E. Shaw et al. (2014) Anton 2: Raising the Bar for Performance and Programmability in a Special-Purpose Molecular Dynamics Supercomputer. in *SC ’14: Proceedings of the International Conference for High Performance Computing, Networking*, Storage and Analysis, pp 41–53.

44. K. Vanommeslaeghe et al., CHARMM general force field: A force field for drug-like molecules compatible with the CHARMM all-atom additive biological force fields. J Comput Chem 31, 671–690 (2010).

45. H. Ke, S. Dass, J. M. Morrisey, M. W. Mather, A. B. Vaidya, The mitochondrial ribosomal protein L13 is critical for the structural and functional integrity of the mitochondrion in Plasmodium falciparum. J Biol Chem 293, 8128–8137 (2018).

46. H. Ke et al., Genetic investigation of tricarboxylic acid metabolism during the Plasmodium falciparum life cycle. Cell Rep 11, 164–174 (2015).

47. L. J. Nkrumah et al., Efficient site-specific integration in Plasmodium falciparum chromosomes mediated by mycobacteriophage Bxb1 integrase. Nat Methods 3, 615–621 (2006).

48. A. M. Lehane et al., Characterization of the ATP4 ion pump in Toxoplasma gondii. J Biol Chem 294, 5720–5734 (2019).

49. S. F. Altschul, W. Gish, W. Miller, E. W. Myers, D. J. Lipman, Basic local alignment search tool. J Mol Biol 215, 403–410 (1990).

50. N. R. Coordinators, Database Resources of the National Center for Biotechnology Information. Nucleic Acids Res 45, D12–D17 (2017).

51. C. Aurrecoechea et al., EuPathDB: the eukaryotic pathogen genomics database resource. Nucleic Acids Res 45, D581–D591 (2017).

52. R. E. Desjardins, C. J. Canfield, J. D. Haynes, J. D. Chulay, Quantitative assessment of antimalarial activity in vitro by a semiautomated microdilution technique. Antimicrob Agents Chemother 16, 710–718 (1979).

