## Supplementary Information for "Mutational Analysis of an Antimalarial Drug Target *Pf*ATP4"

### Supplemental Methods:

#### *Phylogenetic Analysis*

Accession numbers of the sequences chosen for alignment are listed in **Supplementary Table S1**. Sequences from stramenophile and chlorophyte groups were not previously considered in similar phylogenetic analyses. Sequences of P2A ATPases, P2A being among the closest sister clades to P2D (1, 2), were chosen as outgroup sequences. Alignments generated by TCoffee (3), MAFFT (L-INS-i), and MAFFT (G-INS-i) (4) with various parameter settings were examined, and all presented largely similar conserved regions; the final alignment was generated using MAFFT (G-INS-i+homologues). Unambiguously aligned columns were chosen with the assistance of Guidance II (5). The trimmed alignment was subjected to maximum likelihood phylogenetic analysis by PhyML (6) using the optimal substitution model, LG, chosen by SMS (7). Phylogenetic tree output was viewed and arranged for presentation using the Tree Explorer module in the MEGA 6 package (8) and enhanced labeling using Inkscape (http://www.inkscape.org).

#### *Proteomic analysis*

In-gel digestion

Each gel band was subjected to reduction with ten mM DTT for 30 mins at 60 °C, and alkylation with 20 mM iodoacetamide for 45 min at room temperature in the dark and overnight digestion with trypsin (Sequencing grade, Thermo Fisher Scientific) at 37 °C. Peptides were extracted twice with 5% formic acid and 60% acetonitrile and dried under vacuum.

Liquid chromatography-tandem mass spectrometry (LC-MS/MS)

Samples were analyzed using a Q Exactive HF tandem mass spectrometer coupled to a Dionex Ultimate 3000 RLSCnano System (Thermo Scientific). Samples were loaded onto a fused silica trap column Acclaim PepMap-100, 75 μm x 2 cm (Thermo Scientific). After washing for 5 min at five μl/min with 0.1% trifluoroacetic acid, the trap column was brought in line with an analytical column (Nanoease MZ peptide BEH C18, 130 Å, 1.7 μm x 75 μm x 250 mm, Waters) for LC-MS/MS. Peptides were eluted using a segmented linear gradient of 4 to 90% B (A: 0.2% formic acid, B: 0.08% formic acid, 80% acetonitrile): 4–15% B for 5 min, 15-50% B for 50 min, and 50-90% B for 15 min. Mass spectrometry data was acquired using a data-dependent acquisition procedure with a cyclic series of a full scan with a resolution of 120,000 followed by MS/MS (HCD, relative collision energy 27%) of the 20 most intense ions and a dynamic exclusion duration of 20 sec.

Database Search

The peak list of the LC-MS/MS was generated by Thermo Proteome Discoverer (v. 2.1) into MASCOT Generic Format (MGF) and searched against the *3D7* database at PlasmoDB and *Pf*ATP4 specific *Dd2* sequences in addition to a database composed of common lab contaminants using in house version of X!Tandem (9). Search parameters were as follows: fragment mass error: 20 ppm, parent mass error: +/− 7 ppm; fixed modification: carbamidomethylation of cysteine; flexible modifications: Oxidation on Methionine; protease specificity: trypsin (C-terminal R/K unless followed by P), with one miss-cut at preliminary search and five miss-cut during refinement. Only spectra with log e <−2 were included in the final report.

### Supplemental Data

| Source species | Short identifier code | NCBI/EMBL accession number |
|---|---|---|
| *Plasmodium falciparum* | *PLAfa* | XP_024329152.1 |
| *P. gallinaceum* | *PLAga* | XP_028528696.1 |
| *P. vivax* | *PLAvi* | VUZ98172.1 |
| *Babesia bovis*, , | *BABbo* | XP_001610924.1 |
| *Theileria parva* | *THEpa* | XP_766241.1 |
| *Toxoplasma gondii* | *TOXgo* | XP_018635122.1 |
| *Besnoitia besnoiti* | *BESbe* | PFH31327.1 |
| *Eimeria maxima* | *EIMma* | XP_013334395.1 |
| *Cryptosporidium parvum* | *CRYpa* | XP_625857.1 |
| *C. muris* | *CRYmu* | XP_002140372.1 |
| *Vitrella brassicaformis* | *VITbr1* | CEM38216.1 |
| *Chromera velia* | *CHRve* | CEM43005.1 (EMBL) |
| *Vitrella brassicaformis* | *VITbr2* | CEM23227.1 |
| *Symbiodinium microadriaticum* | *SYMmi* | OLQ04979.1 |
| *Breviolum minutum*# | BRE*mi* | symbB1.v1.2.025174 |
| *Micromonas pusilla* | *MICpu* | XP_003062064.1 |
| *Micromonas commoda* | *MICco* | XP_002507786.1 |
| *Aureococcus anophagefferens* | *AURan* | XP_009033039.1 |
| *Aphanomyces invadans* | *APHin* | XP_008873575.1 |
| *Plasmopara halstedii* | *PLAha* | XP_024574184.1 |
| *Phytophthora infestans* | *PHYin* | XP_002899328.1 |
| *Arabidopsis thaliana* (SERCA) | *ARAth_ECA1* | NP_172259.1 |
| *Homo sapiens* (SERCA) | *HOMsa_SERCA2* | NP_733765.1 |
| *Plasmodium falciparum* (SERCA) | *PLAfa_ATP6* | XP_001350994.1 |
| *Physcomitrella patens* | *PHYpa1* | CAD91917.1 |
| *Physcomitrella patens* | *PHYpa2* | *CAD91924.1* |
| *Marchantia polymorpha* | *MARpo* | CAX27437.1 |
| *Riccia fluitans* | *RICfl* | FN691478.1 |
| *Saccharomyces cerevisiae* | *SACce* | NP_010325.1 |
| *Yarrowia lipolytica* | *YARli* | XP_499639.1 |
| *Neurospora crassa* | *NEUcr1* | *CAB65298.1* |
| *Coccidioides posadasii* | *COCpo* | XP_003069007.1 |
| *Neurospora crassa* | *NEUcr3* | XP_962099.1 |
| *Fusarium graminearum* | *FUSgr* | XP_011321377.1 |
| *Schizophyllum commune* | *SCHco* | XP_050201852.1 |
| *Trichophyton rubrum* | *TRIru* | XP_003238625.1 |
| *Allomyces macrogynus* | *ALLma* | KNE72470.1 |
| *Trypanosoma cruzi* | *TRYcr* | XP_817442.1 |
| *Trypanosoma brucei* | *TRYbr* | XP_827683.1 |
| *Leishmania donovani* | *LEIdo* | AAC19126.1 |
| *Leishmania major* | *LEIma* | XP_003722601.1 |
| *Entamoeba invadens* | *ENTin* | XP_004184491.1 |
| *Entamoeba histolytica* | *ENThi* | XP_657556.1 |

#formerly *Symbiodinium minutum) Breviolum minutum* sequence retrieved from the genome project: https://marinegenomics.oist.jp/symb/viewer/info?project_id=21 (10).

**Table S1: Non-abbreviated Version of P2D-type ATPases used for Phylogenetic Analysis with Short Identifier Code and Accession Number**

| | E1P-ADP | E2P | E2P-ATP | E2-ATP |
|---|---|---|---|---|
| **RMSD (whole)** | 6.24 | 7.36 | 7.31 | 6.96 |
| **RMSD (TM region)** | 3.29 | 5.14 | 3.72 | 3.09 |
| **Distance between N and P** | 47.2±1.1 | 26.6±0.7 | 49.5±0.9 | 52.2±1.2 |

**Table S2.** RMSD in Å between the final structures of the various *Pf*ATP4 models from a 115ns MD simulation with the alpha fold structure. The last row includes the center of mass distance in Å between the N and P domains.

| System | Number of Lipids |
|---|---|
| **$Na_2$E1P-ADP** | 321 |
| **E2P** | 510 |
| **E2P-ATP** | 303 |
| **H3E2-ACP** | 312 |

**Table S3.** The total number of lipids used in each of the *Pf*ATP4 systems in MD simulation.

| Oligonucleotide | Primers | Primer Sequence |
|---|---|---|
| **PSC *Pf*ATP4-3Myc; *Pf*ATP4-E409I-3HA** (Na+ Coordination) | Forward Primer | 5'-GTA TCT TCC ATT CCA **ATA** GGT TTA CCT ATG GTT GTT ACT ATC |
| | Reverse Primer | 5'-GAT AGT AAC AAC CAT AGG TAA ACC **TAT** TGG AAT GGA AGA TAC |
| **PSC *Pf*ATP4-3Myc; *Pf*ATP4-E1176I-3HA** (Na+ Coordination) | Forward Primer | 5'-GCT GTT TGG TGT **ATA** ATG CTT AGA GCT TAT ACA GTA AGA |
| | Reverse Primer | 5'-AGC TCT AAG CAT TAT ACA CCA AAC AGC TGA TAT AAA |
| **PSC *Pf*ATP4-3Myc; *Pf*ATP4-D451N-3HA** (Phosphorylation Cycle) | Forward Primer | 5'-TGC TGT TCA GTC ATA TGT TCT **AAT** AAA ACC GGT ACA TTA |
| | Reverse Primer | 5'-TGT CAT TTT TCC TTC AGT TAA TGT ACC GGT TTT **ATT** AGA ACA TAT GAC |
| **PSC *Pf*ATP4-3Myc; *Pf*ATP4-P412T-3HA** (Resistance to Spiroindolones and DHIQ's) | Forward Primer | 5'-CTTCCATTCCAGAAGGTTTAACTATGGTTGTTACTATCACC- |
| | Reverse Primer | 5'-GGTGATAGTAACAACCATAGTTAAACCTTCTGGAATGGAAG |
| ***PfATP4-3Myc; *Pf*ATP4-V178I+G223R-3HA** (Resistance to Pyrazoleamides and Spiroindolones) *gDNA was PCR amplified from an NF54 parasite line exposed to Cmpd2-1 containing a V178I mutation. This product was used with the primers in this chart to create the V178I+G223R double mutant | Forward Primer | 5'-GAAAAATCATCAGGTGATGCTATACGAAAATTAGCTGAAATGGCTTCAC |
| | Reverse Primer | 5'-GTGAAGCCATTTCAGCTAATTTTCGTATAGCATCACCTGATGATTTTTC |
| **Integration Primers (Nkrumah, Nat Methods, 2006)** | Forward Primer | 5'-CATTTGAATTATTGCTCAACGCT |
| | Reverse Primer | 5'-GATAGCGATTTTTTTTACTGTCTG |
| **Primers to sequence ectopic RL2 *Pf*ATP4-3HA from *Plasmodium falciparum* genomic DNA** | Forward Primer | 5'-TAA TAT TAT ACA ATA TAC CTA GGT GAT AAA TGA GTT CTC |
| | Reverse Primer | 5'-TGA CAC TAT AGA ATA CTC AAG CTT GC |
| **Primers to sequence ectopic CAM *Pf*ATP4-3HA from *Plasmodium falciparum* genomic DNA** | Forward Primer | 5'-ATA ATA ATA AAT ACC TAA TAG AAA TAT ATC ACC TAG G |
| | Reverse Primer | 5'-TGA CAC TAT AGA ATA CTC AAG CTT GC |

**Table S4: List of Oligonucleotides** used for (1) generation of merodiploid, mutant lines, (2) checking proper integration of ectopic PfATP4, and (3) PCR amplification of ectopic *Pf*ATP4 from parasite genomic DNA.

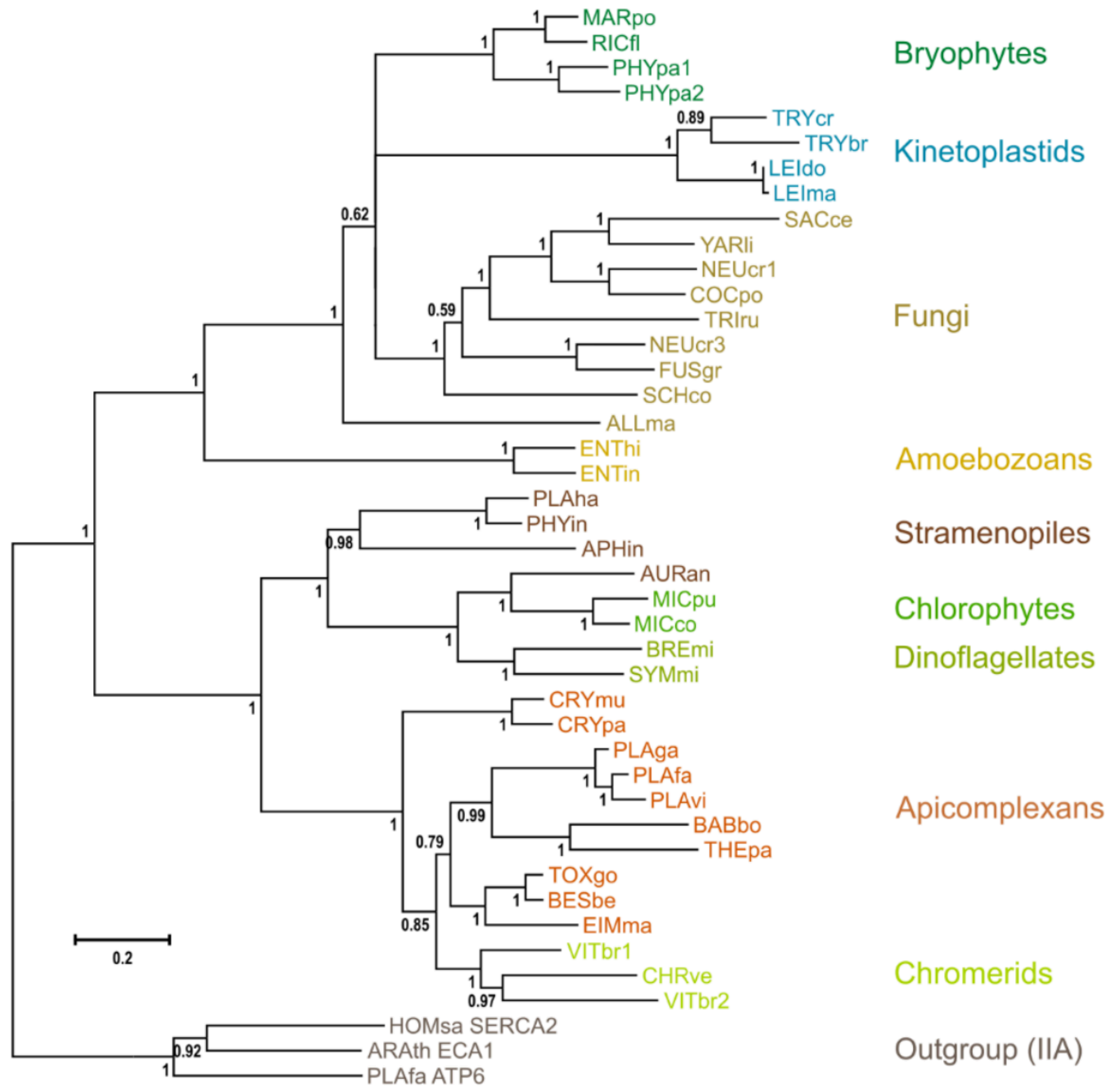


**Figure S1: Rectangular Phylogram of P2D-type ATPases.** Phylogenetic tree of type P2D ATPases with bootstrap values indicated. Sequences included and used in the figure are identified using the short identifier (Uniprot style). Accession numbers and full versions of the genus and species are given in **Table S1** with highlighted colors.

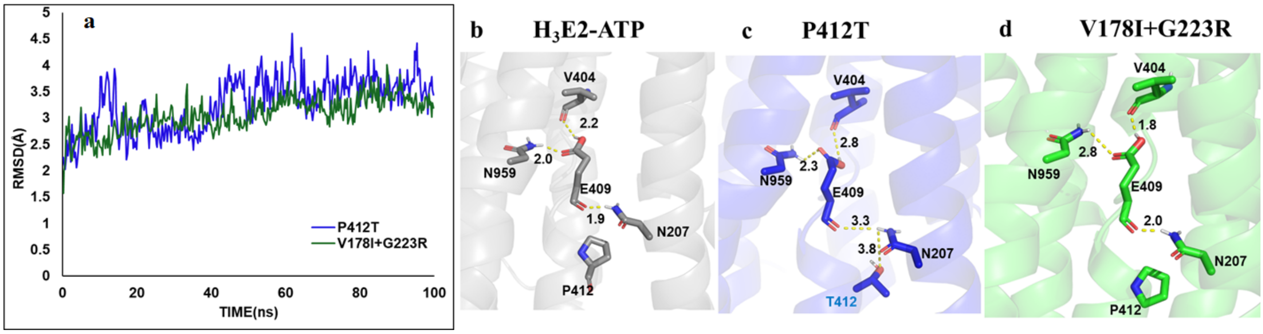


**Figure S2: Mutants P412T and V178I+G223R significantly change the H-bonding pattern at the $Na^+$ binding site** (a) trajectory traces from MD simulations shows both mutant models are stably equilibrated throughout the simulation; three dimensional models of *Pf*ATP4 models are rendered as grey ribbons from wild type (b), blue ribbons for the P412T mutant (c), and green ribbons for the V178I+G223R mutant (d). The H-bond interactions of E409 with N207, V404 and N959 in each model are represented as yellow dotted lines and the distances are measured in Å. Participating residues are shown as licorice sticks and labeled.

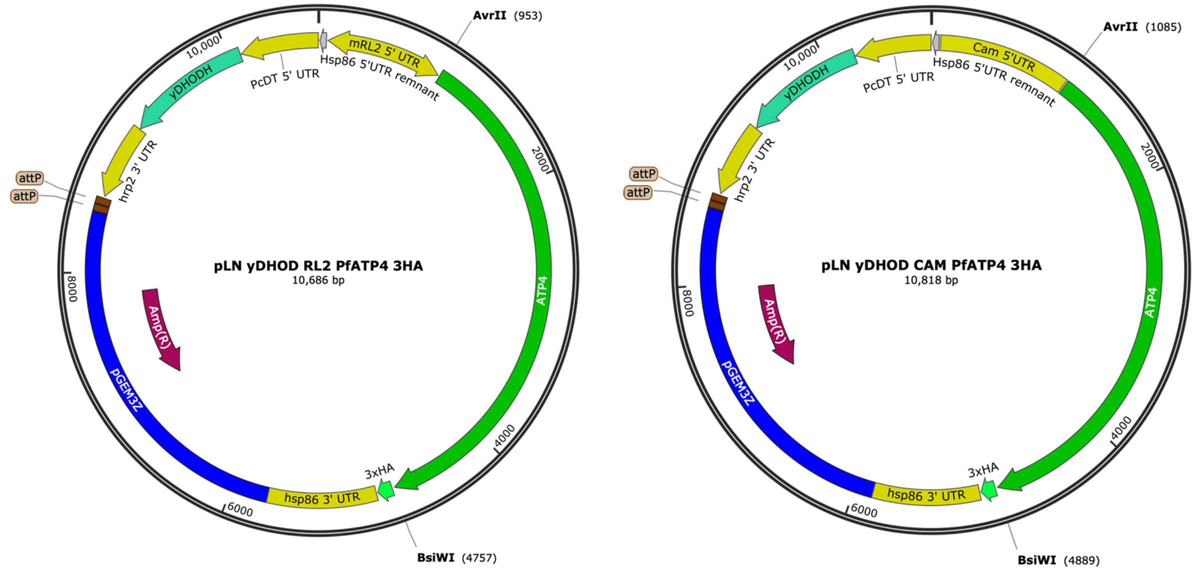


**Figure S3: Plasmid Maps of pLN vectors** used for transfection of NF54:*attb Pf*ATP4:3xMyc parasites to generate merodiploid NF54*attB Pf*ATP4:3xMyc *Pf*ATP4$_{(Mut)}$:3xHA parasites. Ectopic expression of *Pf*ATP4 in *Pf*ATP4$_{WT}$, *Pf*ATP4$_{E409I}$, *Pf*ATP4$_{E1176}$, and *Pf*ATP4$_{D541N}$ merodiploid parasite lines was regulated by the RL2 promoter whereas ectopic *Pf*ATP4 expression in *Pf*ATP4$_{P412T}$, and *Pf*ATP4$_{V178I+G223R}$ parasites lines was controlled by the CAM promoter.

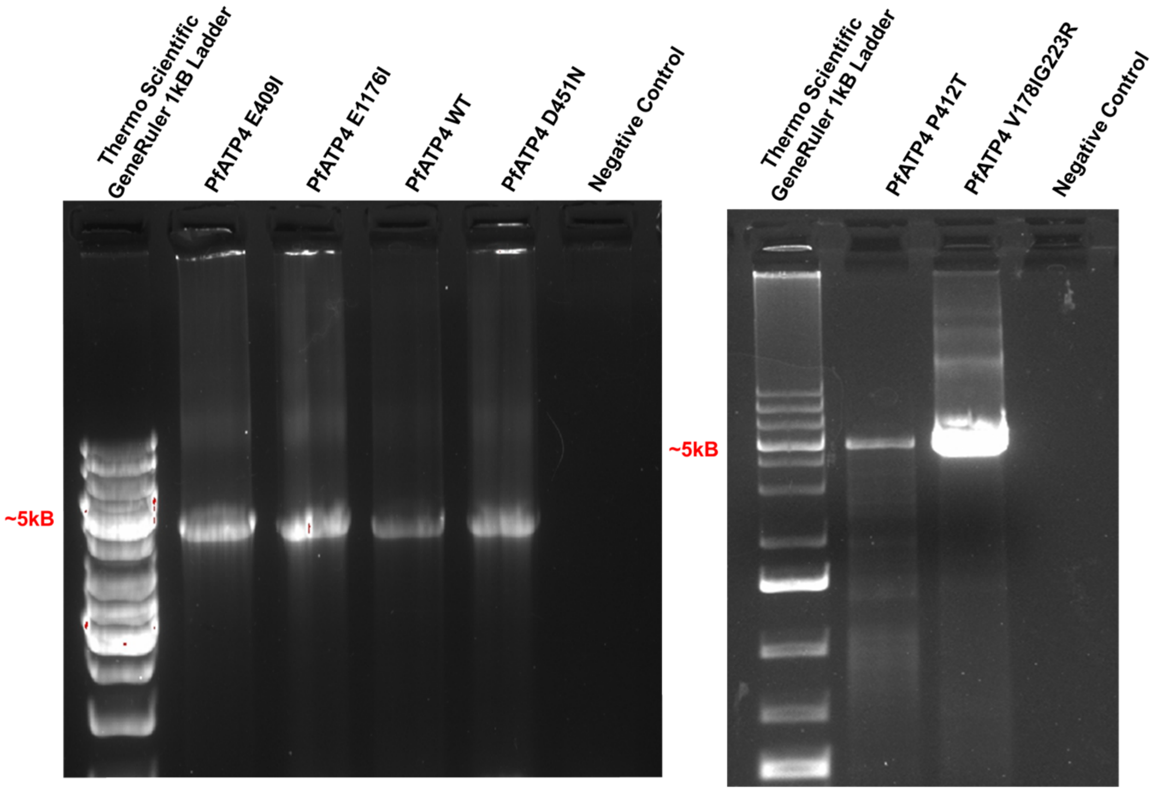


**Figure S4**: **Confirmation of Integration at *attB* site.** Integration of ectopic *Pf*ATP4$_{mut}$ (all lines) to the *attB* site.

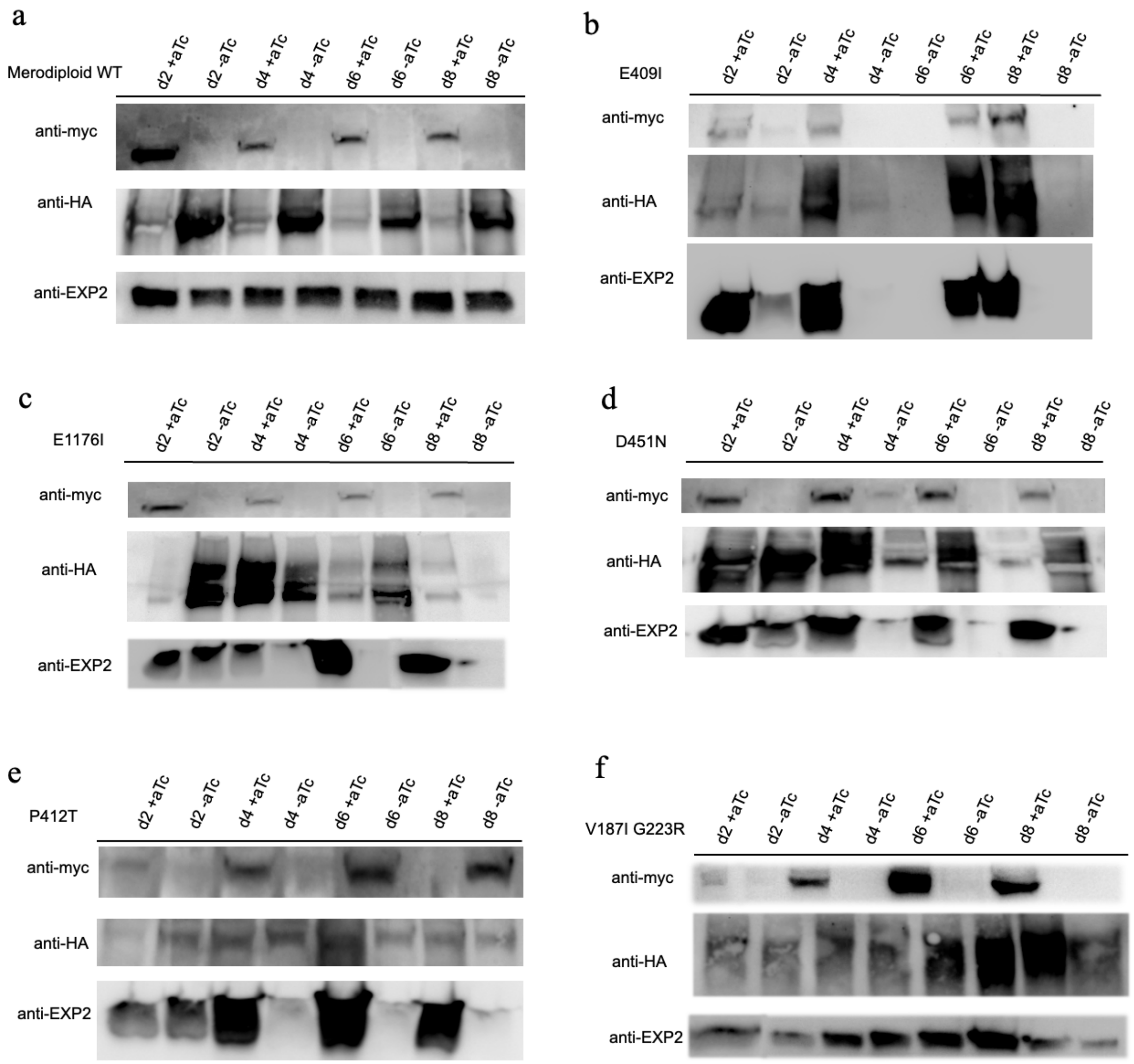


**Figure S5: Myc and HAWestern Blots of Merodiploid Parasite Lines.** Western Blots to confirm knockdown of endogenous P*f*ATP4:3xMyc in the absence of aTc and to confirm proper expression of P*f*ATP4$_{mut}$:3xHA in the presence and absence of aTc.

**Supplementary References**


1. A. Rodriguez-Navarro, B. Benito, Sodium or potassium efflux ATPase a fungal, bryophyte, and protozoal ATPase. *Biochim Biophys Acta* **1798**, 1841-1853 (2010).
2. A. M. Lehane *et al.*, Characterization of the ATP4 ion pump in Toxoplasma gondii. *J Biol Chem* **294**, 5720-5734 (2019).
3. P. Di Tommaso *et al.*, T-Coffee: a web server for the multiple sequence alignment of protein and RNA sequences using structural information and homology extension. *Nucleic Acids Res* **39**, W13-17 (2011).
4. K. Katoh, K. Kuma, H. Toh, T. Miyata, MAFFT version 5: improvement in accuracy of multiple sequence alignment. *Nucleic Acids Res* **33**, 511-518 (2005).
5. I. Sela, H. Ashkenazy, K. Katoh, T. Pupko, GUIDANCE2: accurate detection of unreliable alignment regions accounting for the uncertainty of multiple parameters. *Nucleic Acids Res* **43**, W7-14 (2015).
6. S. Guindon *et al.*, New algorithms and methods to estimate maximum-likelihood phylogenies: assessing the performance of PhyML 3.0. *Syst Biol* **59**, 307-321 (2010).
7. V. Lefort, J. E. Longueville, O. Gascuel, SMS: Smart Model Selection in PhyML. *Mol Biol Evol* **34**, 2422-2424 (2017).
8. K. Tamura, G. Stecher, D. Peterson, A. Filipski, S. Kumar, MEGA6: Molecular Evolutionary Genetics Analysis version 6.0. *Mol Biol Evol* **30**, 2725-2729 (2013).
9. R. Craig, J. P. Cortens, R. C. Beavis, Open source system for analyzing, validating, and storing protein identification data. *J Proteome Res* **3**, 1234-1242 (2004).
